# Multiple blood feeding bouts in mosquitoes allow for prolonged survival and are predicted to increase viral transmission during drought

**DOI:** 10.1101/2024.05.28.595907

**Authors:** Christopher J. Holmes, Souvik Chakraborty, Oluwaseun M. Ajayi, Melissa R. Uhran, Ronja Frigard, Crystal L. Stacey, Emily E. Susanto, Shyh-Chi Chen, Jason L. Rasgon, Matthew DeGennaro, Yanyu Xiao, Joshua B. Benoit

**Affiliations:** Department of Biological Sciences, University of Cincinnati, Cincinnati, OH; Department of Entomology, Center for Infectious Disease Dynamics and Huck Institutes for Life Sciences, Pennsylvania State University, University Park, Pennsylvania, PA; Department of Biological Sciences and Biomolecular Sciences Institute, Florida International University, Miami, FL; Department of Mathematical Sciences, University of Cincinnati, Cincinnati, OH

**Keywords:** Mosquito refeeding, vectorial capacity, disease transmission modeling, host-sensing, activity, sleep, climate change

## Abstract

Survival through periods of drought is critical for mosquitoes to reside in semi-arid regions with humans. Dry conditions increase blood feeding propensity in mosquitoes, but it is unknown if dehydration-induced bloodmeals increase feeding beyond what is necessary for reproduction. Following a bloodmeal, prolonged exposure to dry conditions increased secondary blood feeding in mosquitoes by nearly two-fold, and chronic blood feeding allowed mosquitoes to survive up to twenty days without access to water sources. This refeeding did not alter the number of eggs generated, suggesting this refeeding is for hydration and nutrient replenishment. Exposure to desiccating conditions following a bloodmeal resulted in increased activity, decreased sleep levels, and prompted a return of CO_2_ sensing before egg deposition. Increased blood feeding during the vitellogenic stage and higher survival during dry periods are predicted to increase pathogen transmission, allowing for a rapid rebound in mosquito populations when more favorable conditions return. This explains the elevated levels of specific arbovirus cases in association with periodic dry conditions and warrants further consideration as climate change progresses. Overall, these results solidify our understanding of the role of dry periods on mosquito blood feeding and how mosquito dehydration contributes to vectorial capacity and disease transmission dynamics

**Significance statement:** Bouts of dehydration yield substantial changes to insects’ physiology and behavior. Mosquitoes are exceptionally prone to dehydration due to high water loss rates, but few integrative studies have examined the comprehensive impact of drought conditions on mosquitoes. Here, we demonstrate that dry conditions lead to multiple blood feeding events, allowing mosquitoes to survive dry periods. This repeated blood feeding is associated with higher activity and an early return of attraction to vertebrate hosts. Increased dry season survival and more frequent blood feeding are predicted to yield higher transmission of mosquito-borne viruses. This suggests that a higher prevalence of drought associated with climate change will have varying impacts on mosquito-borne diseases.

## Introduction

Blood feeding by female mosquitoes primarily functions as a means for reproduction (1), resulting in the spread of mosquito-borne pathogens. When an infected female mosquito feeds on an uninfected host, a transfer of pathogens is possible. While considerable research has been conducted on pathogen transmission in mosquitoes, more research is needed to examine how environmental factors impact mosquito feeding (2–4) and refeeding in the context of vectorial capacity. Recent studies have uncovered interesting developments involving these factors, such as thermal changes, their role in pathogen transmission, and how drought impacts blood feeding (2–4). These ecophysiological studies have increased our understanding of the direct effects of environmental factors on mosquito physiology and behavior. Still, studies on the impact of dry periods on mosquito biology remain limited compared to the focus on thermal effects (5–7).

Drought exposure and dehydration significantly impact mosquitoes, shifting population levels, altering feeding propensity, and increasing pathogen transmission (5–8). Of interest, dry periods have been linked to increased transmission of arboviruses (9–11), which is unexpected as periods with reduced water availability have been associated with lower mosquito populations (12, 13). This increased viral transmission has been linked to specific environmental factors, which allows for increased mosquito blood feeding and survival rates. For example, culverts become stagnant, altering mosquito-predator interactions and enabling mosquito growth (5, 6, 14). When mosquitoes are blood-fed before dehydration, individuals show an increased survival time compared to their non-blood-fed counterparts, likely due to higher water content at the start of dehydration exposure (6). These findings underscore the critical function of blood feeding for the replenishment of water content in dehydrated mosquitoes (5, 6, 15). However, the extent of water replenishment by blood feeding during short dry conditions (hours to days), that can extend into a drought (days to weeks), has yet to be examined. Importantly, increased periods of drought are expected to increase with climate change (16), indicating that understanding this process will be critical for mosquito biology and disease transmission in future scenarios.

Interactions between multiple bloodmeals during a single reproductive cycle and the resulting hydration status of mosquitoes have also not been examined. This is surprising as refeeding has been implicated in many other biological changes that occur following dehydration, including fecundity, nutritional reserve supplementation, and virus dissemination within the mosquito (17, 18). Notably, a recent study on refeeding in *Aedes aegypti* established higher rates of viral dissemination when a second (uninfected) bloodmeal was allowed within the extrinsic incubation period (17). These refeeding events produced disturbances and microperforations in the basal lamina, which could increase viral particles escaping from the midgut, resulting in increased viral transmission potential (17). Combining increased pathogen dissemination following a second bloodmeal (17) with the propensity of mosquitoes to blood feed when dehydrated (5, 6), mosquitoes exposed to dry conditions may exhibit increased blood feeding and viral transmission rates during periods when refeeding is typically unexpected.

Multiple bloodmeals could be common within mosquito species to promote egg production, where up to 10-40% of mosquitoes will ingest a second bloodmeal before the previous blood has been digested (19–24). Ordinarily, a combination of mechanisms, predominantly regulated by hormonal cues, suppress additional blood feeding (25, 26), but when feeding to repletion is interrupted, the signals that prevent a secondary bloodmeal can be suppressed (20, 24, 27). Another such mechanism is the acquisition of nutrients through a secondary bloodmeal to supplement poor larval nutrition levels, which can be necessary for oogenesis (20, 24, 27). Even though multiple blood feedings are observed, a general reduction in blood-fed females in host mimic traps suggests that host attraction is likely to be reduced even if a second meal is possible (28–30). This reduction in post-blood feeding attraction is supported by specific molecular changes in the mosquito neurotranscriptome that reduce responsiveness to host cues (31). An overlooked aspect of this interaction is that the bloodmeal can represent a substantial source of hydration (6), allowing immediate replenishment of water stores in dehydrated mosquitoes. After the bloodmeal has been processed and excess fluid from the blood has been expelled, a blood-fed female requires 3-4 days to process the blood under preferred conditions (26, 32). During this time, a female typically resides in an environment that prevents dehydration until egg development is complete (33, 34). If a suitable refuge is not found, dehydration is likely to occur, and increased water ingestion from blood, nectar, or free water sources will be required (5, 35, 36). A common assumption is that mosquitoes feed once and produce eggs before a second bloodmeal is consumed (20, 26), but multiple blood feedings have been previously observed in both *Aedes spp.* and *Anopheles spp.* (19, 21, 23). Significantly, prolonged dry periods will reduce free water and the water content of nectar sources (5, 37), suggesting that blood from vertebrates may be a more reliable source of water, especially for anthropophilic and endophilic mosquito species (5, 15, 35, 36).

Here, we establish the role of dehydrating conditions on the refeeding propensity of mosquitoes. Specifically, we assess the impact of drought-induced refeeding on survival and reproduction, behavioral changes that occur to increase blood feeding, and how these changes can allow for mosquito survival and pathogen transmission during dry periods. Briefly, these studies revealed a drastic increase in refeeding during dry periods associated with a resumption of attraction to CO_2_ before the end of a gonotrophic cycle, allowing mosquitoes to survive as adults through periods with low water availability. Transmission and population modeling suggests that this increase in secondary feeding may directly underlie mosquito survival through periods of low water availability, and ultimately result in higher drought-induced viral transmission.

## Results

### Dry conditions result in substantial refeeding that allows extended survival

Mosquitoes allowed to blood feed and then held under dehydrating conditions (30-40% RH without access to water sources) had substantial increases in blood feeding during vitellogenesis (Fig. 1). Specifically, one day of post-blood feeding dehydration had a minimal impact on refeeding, but 48 hours under these dry conditions yielded increased feeding under small, medium, and large cage assays (Fig. 1, observed for *An. Stephensi* - tested in small cages only). An average of 2.5 feeding events occurred in medium-sized cages, and 2.0 feeding events occurred in large cages, when mosquitoes were held under dry conditions (Fig. 1D-E, Fig. S1). Under these conditions, most, if not all, mosquitoes would undergo at least one, if not two, additional feedings before egg deposition in a single gonotrophic cycle (Fig. 1). Of importance, these refeeding events allowed the mosquitoes to survive through periods of no water for up to twenty days (Fig. 2B). When allowed to deposit eggs after a prolonged period of chronic blood feeding, there was a reduction in eggs deposited compared to those that were allowed to deposit only four days after a bloodmeal (Fig. 2B). This suggests that prolonged retention will allow eggs to remain viable, a finding that has been previously observed when oviposition sites are not available (38). Overall, these studies indicate that dry conditions are likely to result in the ingestion of multiple bloodmeals during each gonotrophic cycle, allowing females to survive for extended dry periods.

**Figure 1.**
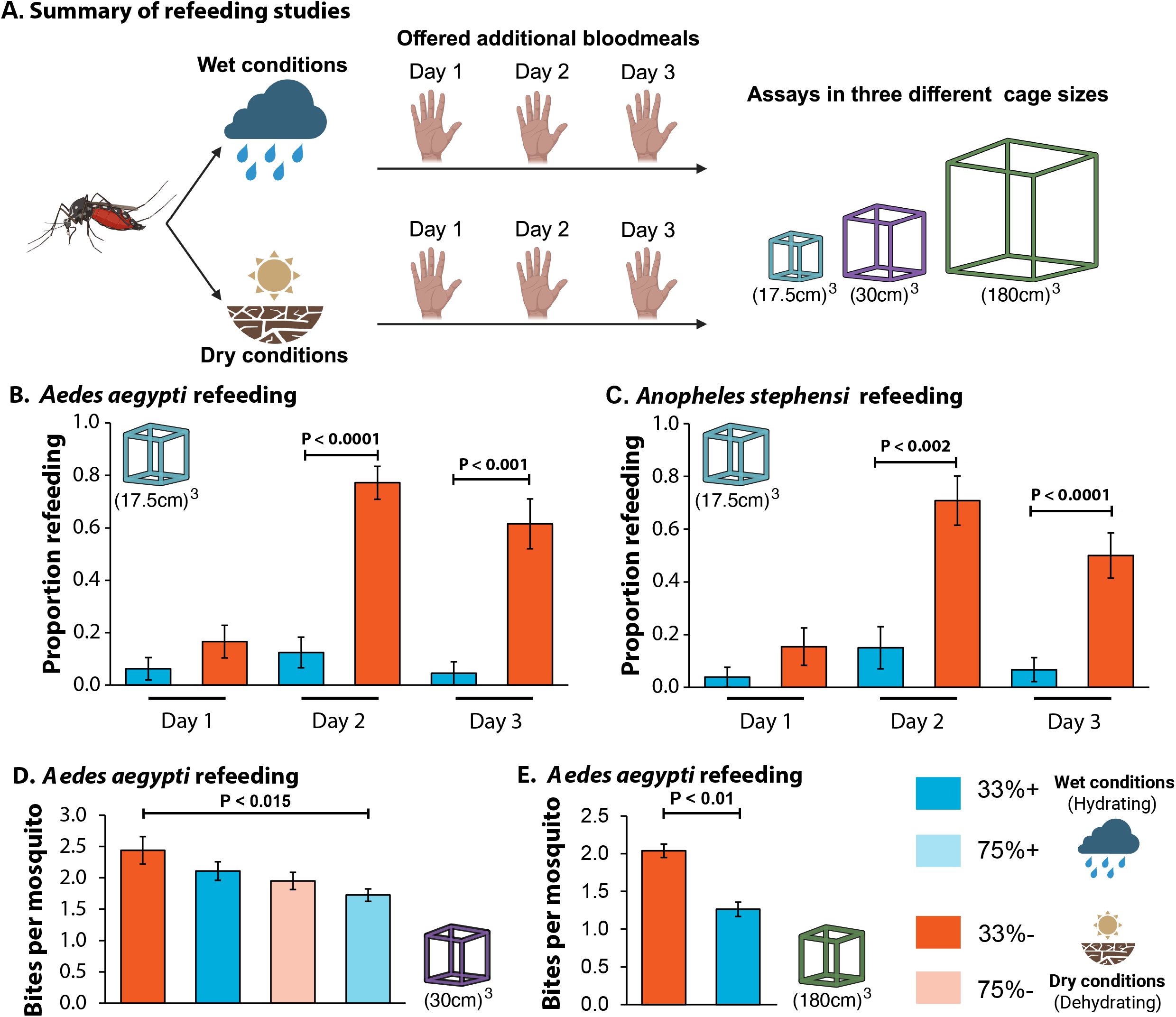
Dehydration induces refeeding in mosquitoes. A. Summary of the studies with mosquitoes held under wet (hydrating) and dry (dehydrating) conditions. B. and C. Small cages (10 x 10 x 10 cm, single mosquitoes, N=15-25) for *Aedes aegypti* and *Anopheles stephensi*, D. Medium cages (30 x 30 x 30 cm, 10 mosquitoes, N=4), and E. Large cages (1.8 x 1.8 x 1.8 m, N=8, ten mosquitoes per cage). Dry conditions increased refeeding for both *Ae. aegypti* and *An. stephensi* (Pairwise Chi-Squared test, ANOVA with Tukey HSD, and Student’s t-test, P < 0.05) under small, medium, and large cage sizes, respectively.

**Figure 2.**
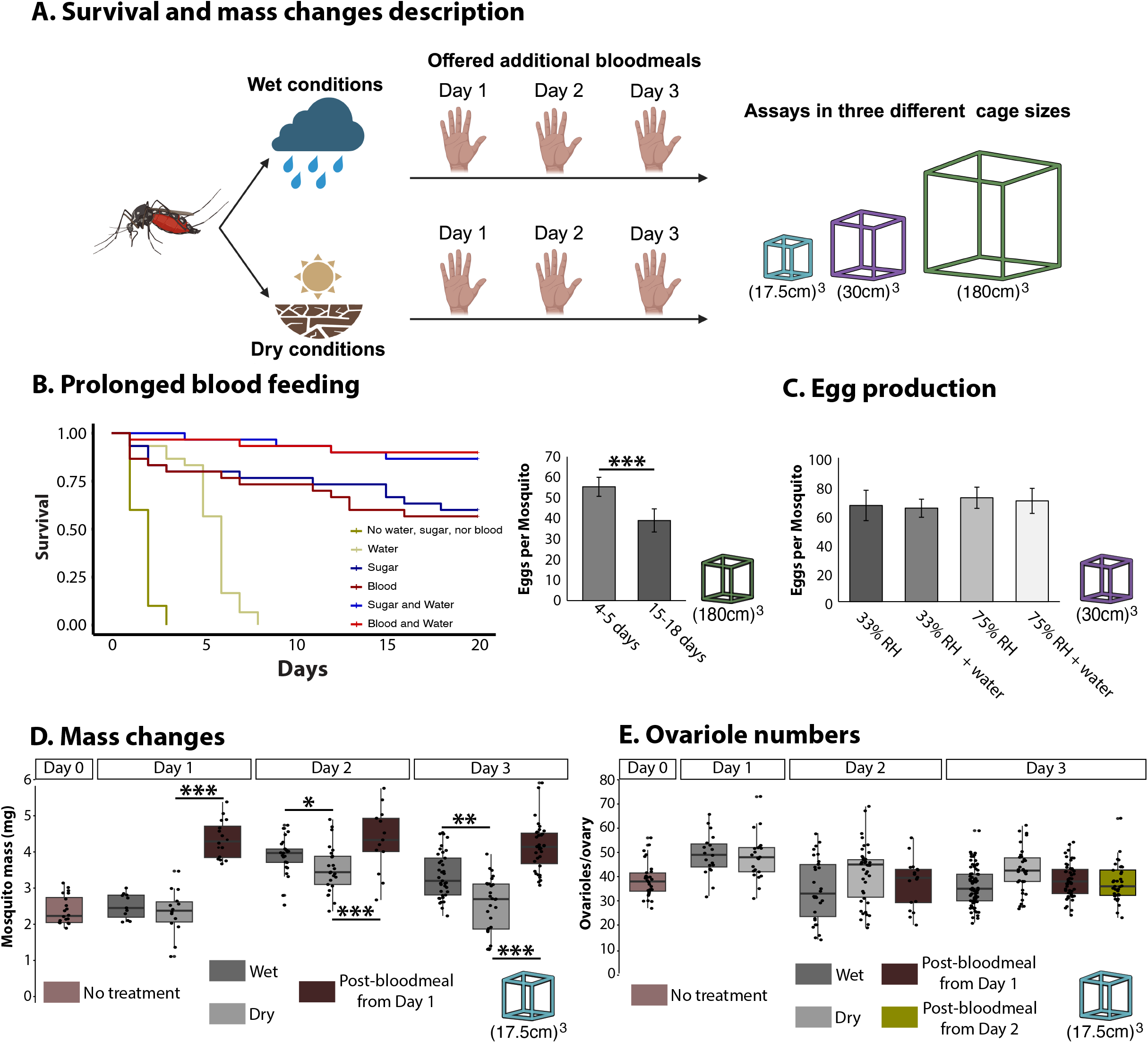
Mosquito mass changes and reproductive output following the secondary bloodmeal. A. Summary of the studies with mosquitoes held under wet (hydrating) and dry (dehydrating) conditions. B. Left - Survival of *Aedes aegypti* under dry and wet conditions with and without access to water, sucrose solution, and blood (30 x 30 x 30 cm, 10 mosquitoes, N=3), Right - Egg production following prolonged periods of refeeding (30 x 30 x 30 cm, 10 mosquitoes, N=3). C. Egg production by *Ae. aegypti* in medium cages (30 x 30 x 30 cm, 10 mosquitoes, N=4). D and E. Mass and ovariole changes measured in small cages (10 x 10 x 10 cm, single mosquitoes, N =15-25) for *Ae. aegypti*, and each group was held under wet or dry conditions following feeding. Prolonged refeeding reduced the number of eggs oviposited, and bloodmeals resulted in consistently higher masses in *Ae. aegypti* (ANOVA with Tukey HSD, and Student’s t-test, *, P < 0.05; **, P < 0.01 ***, P < 0.001).

### Increased blood feeding does not increase egg production

As increased feeding is typically associated with a higher egg output, we evaluated if increased blood feeding under dehydrating conditions resulted in the generation of more eggs (Fig. 2). When single mosquitoes were evaluated, oocyte numbers were not shifted following an additional bloodmeal (Fig. 2E), and in medium cage assays, the same number of eggs per mosquito were deposited, regardless of exposure to dry conditions (Fig. 2C). However, the eggs per bloodmeal were reduced, indicating decreased use of blood for vitellogenesis (Fig. S2B). Oocyte size did not vary based on the number of bloodmeals when individual mosquitoes were examined (Fig. S2E-F). Even one day after a secondary bloodmeal, the oocyte size and number did not vary compared to the control group (Figs. 2E, S2F). These studies suggest that secondary bloodmeals during dry periods are primarily used to increase water content and survival in lieu of reproductive output.

To assess the replenishment of water stores, we examined mass changes following a bloodmeal (Fig. 2D). A significant loss of mass occurred when mosquitoes were held under dry conditions, featuring a 30-40% decline in mass following a bloodmeal after two days for *Ae. aegypti* (Fig. 2). When held in stable conditions, a decline was noted, but mosquitoes retained nearly 0.8 mg more of mass compared to the dehydrated controls after two days under dehydration. Ingestion of a second bloodmeal allowed the mosquitoes to immediately replenish mass up to the amount after the first bloodmeal. As these changes are predominantly due to water loss, prolonged exposure to dry conditions following a bloodmeal will yield mosquitoes that lose nearly half of their water content within two days if unable to rest in stable, relatively humid areas after a bloodmeal. The water content can be replenished if a mosquito obtains a second bloodmeal. Each period of bloodmeal-associated hydration allows survival for another two days under dry conditions until a water source can be located to ingest free water and deposit eggs. Along with these studies on *Ae. aegypti*, similar effects were noted for *An. stephensi* (Fig. S2C-E).

### Dry conditions increase activity and prompt an early return of carbon dioxide **sensing.**

As dry conditions prompted a substantial increase in secondary blood feeding, we examined how activity and behavior of *Ae*. *aegypti* shifted following a bloodmeal with and without access to water sources (Fig. 3). When activity levels were assessed, a general suppression of activity occurred until three days after a bloodmeal, where there was a significant increase in activity of mosquitoes held under dry conditions (Fig. 3B-C). However, the movement for both groups was still considerably lower than the non-blood fed group (Fig. 3B). As activity increased, we showed that sleep decreased in the dehydrated group (Fig. D). This suggests an early increase in activity and decrease in sleep after a bloodmeal when females do not rest in an environment with sufficient relative humidity to maintain water balance.

**Figure 3.**
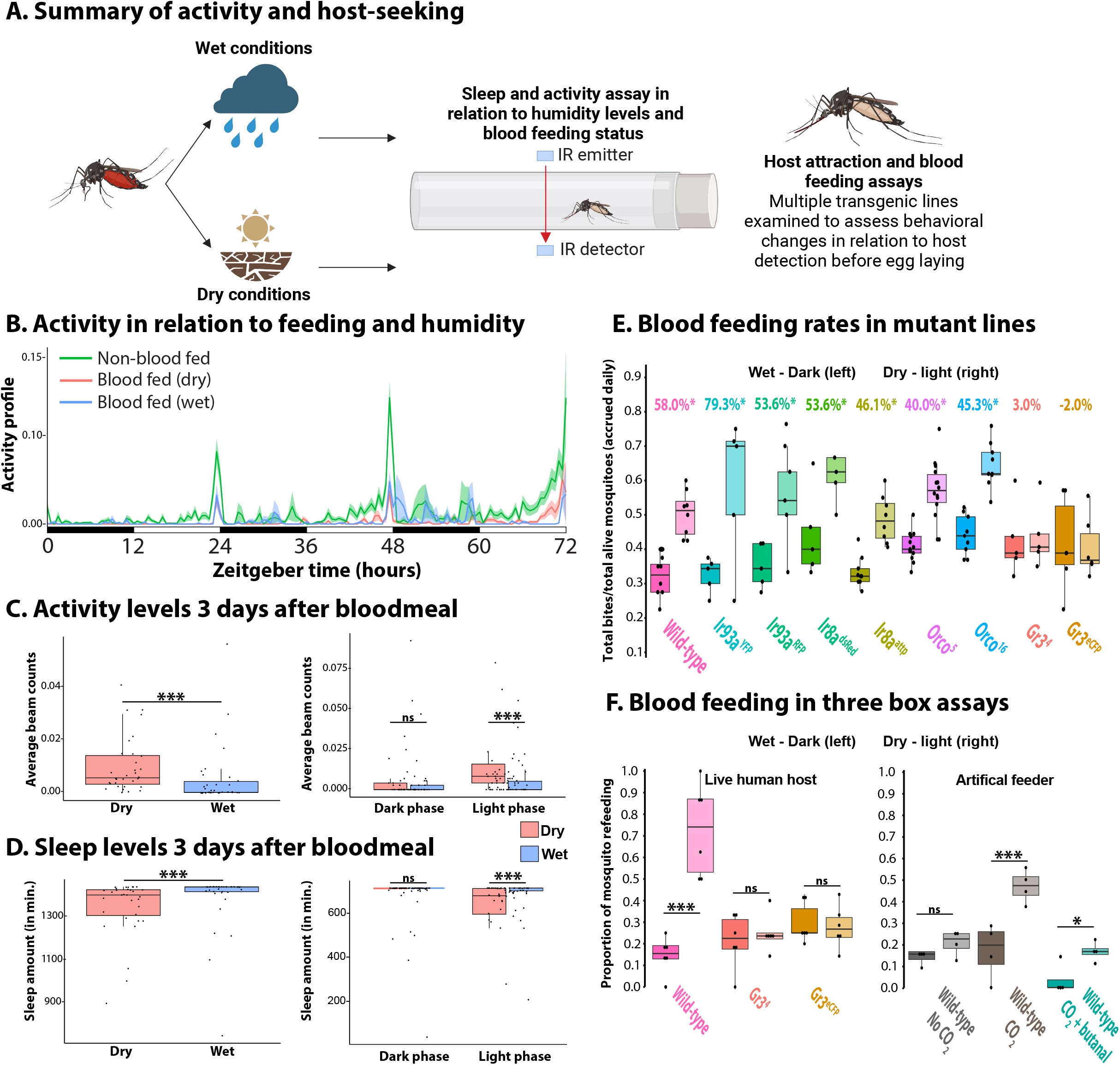
Shifts in activity and impact of impaired host sensing on dehydration-induced refeeding. A. Summary of the studies with mosquitoes held under wet (hydrating) and dry (dehydrating) conditions. B-D. Activity and sleep profiles in blood-fed *Aedes aegypti* when held under dehydrating conditions (33% +/- 5% RH) and hydrating conditions (75% +/- 5% RH with access to water) compared to non-blood-fed mosquitoes with access to 10% sucrose (N=32-48). Dry conditions prompted increased activity and reduced sleep on day three following a bloodmeal, but prior to oviposition (differences driven by light-phase, Wilcoxon rank sum test, P < 0.001). E. Use of *Ae. aegypti* mutant lines to assess host feeding (N=5-14). All mutant lines showed increased blood feeding under dehydration (ANOVA with Tukey HSD, P < 0.05), except for mutant lines with defects in CO_2_ (both GR3 mutant alleles), percent values indicate mean biting differences between dry and wet conditions of each mosquito line. F. Utilization of three-box choice assays to directly examine the role of CO_2_ detection on multiple feeding attempts using both a live host and an artificial feeder (Hemotek). A lack of CO_2_ with artificial feeding or impaired sensing of CO_2_ (Gr3 mutants with a live host or butanal treatment with artificial feeding) leads to a reduction in dehydration-induced refeeding (Pairwise Chi-Squared test, ***, P < 0.0001; *, P < 0.05).

Increases in activity and host feeding suggest there may be a reconstitution of host-sensing, which we examined through the utilization of mutant mosquitoes with impaired or altered host or humidity detection (39–43). In general, most of the mutant strains showed a similar response to the control lines, both in the time it took the mosquitoes to feed on the host (Fig. S3), and in the amount that dehydration increased blood feeding (Fig. 3). The lone exception was Gr3 mutants with impaired CO_2_ sensing, which did not show an increase in blood feeding during dehydration (Fig. 3E). To confirm if increased CO_2_ sensing was associated with dehydration, we performed host feeding assays with and without the presence of CO_2_, which showed increased refeeding on an artificial host when CO_2_ was present (Fig. 3F). A secondary experiment was conducted where mosquitoes were exposed to butanal to reduce response to CO_2_ (44), which confirmed that increased CO_2_ sensing is a significant factor associated with dehydration-induced refeeding (Fig. 3E-F). These studies highlight that dehydration leads to mosquito sensory and behavioral changes that increase blood feeding, even within a single gonotrophic cycle.

### Increased blood feeding and survival will shift viral transmission and mosquito populations

Previous studies have shown that dry conditions increase the transmission of specific viruses (9, 10) and impact viral dissemination in mosquitoes (7, 45). We built upon these observations with our study to model how multiple refeeding events may impact vectorial capacity, mosquito survival, and viral transmission, details are provided in the Supplemental modeling. The multiple feeding events observed under dry conditions are predicted to constitute a 2-3 fold increase in vectorial capacity for mosquitoes (Fig. 4A). When the population growth rates were assessed for *Ae. aegypti* (Fig. 4, Supplemental modeling), the capability of *Ae. aegypti* females to remain viable for extended periods (46). With the extended viability, *Ae. aegypti* can maintain higher populations durign dry periods that can rapidly increase when conditions are more favorable for egg laying and larval growth (Fig. 4B, C). The increased feeding events, and the resultant increase in survival, during dry periods lead to predicted increases in the transmission of Zika by *Ae. aegypti* (Fig. 4D, E). Similar results were obtained for *C. pipiens* in relation to population growth and for West Nile virus transmission (Supplemental modeling).

**Figure 4.**
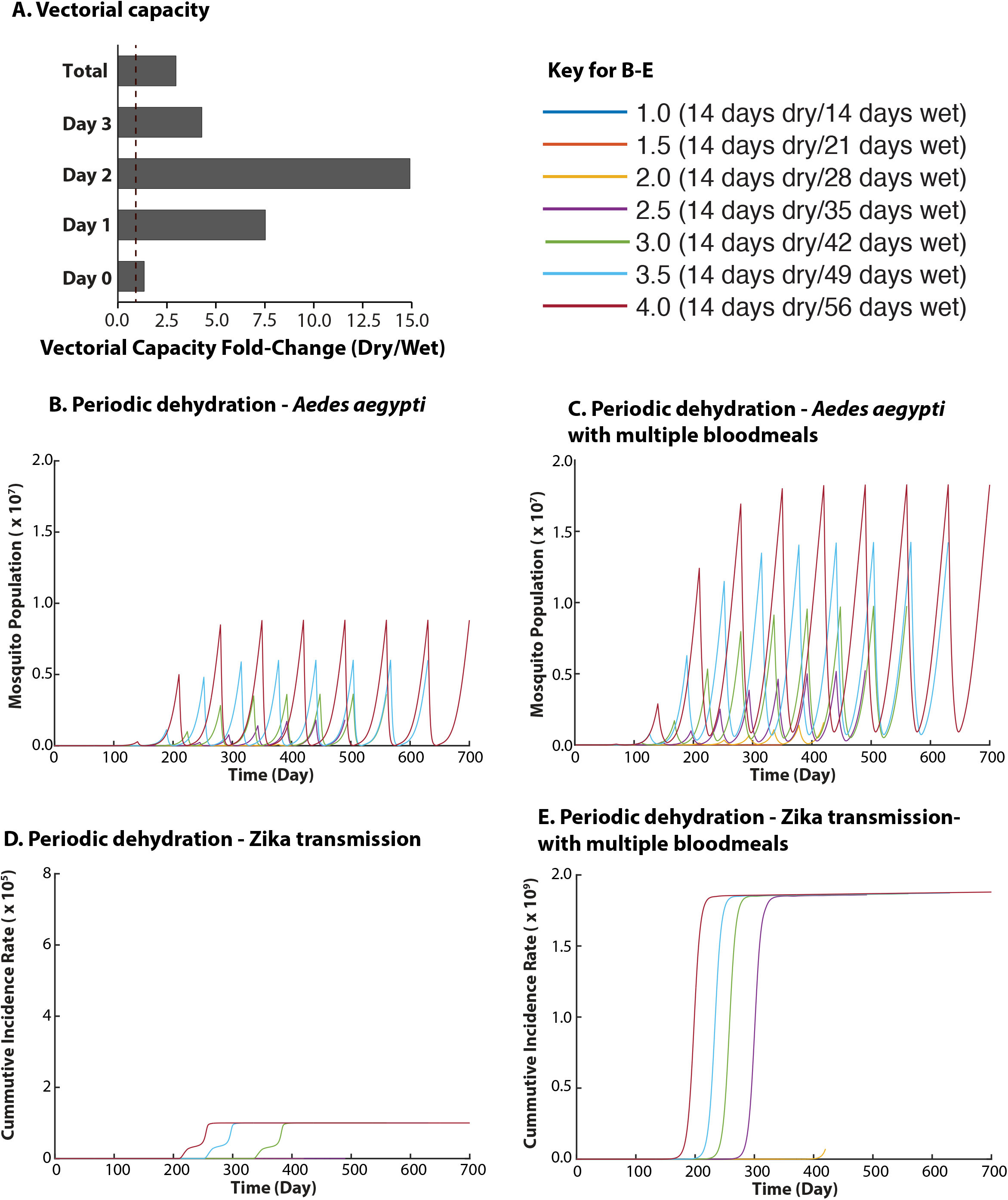
Vectorial capacity and mosquito population growth are increased by multiple bloodmeals. A. Vectorial capacity was determined according to previous studies (6, 70) with modified parameters based on this study. Each day represents post-blood feeding time, with day 0 indicating the fold-change immediately after the initial bloodfeeding. B-F. *Aedes aegypti* population and Zika transmission dynamics in simulated environments, each including dry periods of 14 days with various wet period lengths both without (B) and with access to multiple bloodmeals (C). In D., cumulative incidence rates for Zika virus per 100,000 humans were determined using population estimates from mosquitoes that did not take multiple bloodmeals (shown in B). In E., cumulative incidence rates for Zika virus were determined using population estimates from mosquitoes imbibing multiple bloodmeals (shown in C). Details on modeling are provided in the supplemental materials.

## Discussion

This study established that prolonged exposure to low humidity conditions is a major factor in multiple blood feeding events. Therefore, we expect mosquitoes experiencing water loss will frequently refeed to compensate for lost water when provided ample opportunities. If dehydration persists for an extended period, mosquitoes can prolong survival under desiccating conditions by repeatedly refeeding with no apparent detriments to their behavior or physiology, other than reduced egg production. Our findings indicate that this increase in blood feeding is associated with a return of increased activity and CO_2_ sensing. Modeling based on increased blood feeding rates for rehydration (this study; (5)), increased viral transmission from multiple bloodmeals (17), and prolonged survival by refeeding (this study), indicates that this dehydration-induced refeeding will lead to substantial increases in viral transmission. This provides a key mechanism explaining previous observations that viral transmission increases during dry periods (9, 10).

Although water availability may be limited during dry periods, as floral nectar and liquid water sources are reduced (5, 37), we have previously shown that dry periods will increase mosquito blood feeding propensity (5, 6). We now expand these findings to show substantial increases to blood feeding rates within a single gonotrophic cycle. Previously, multiple blood feeding events have been attributed to an interruption in feeding or a lack of nutrient reserves obtained during larval development (19, 20). In this study, we expand our understanding of refeeding to indicate that dry conditions can be a major underlying factor in this process. As both *An. stephensi* and *Ae. aegypti* have highly anthropophilic tendencies, there could be specific fitness advantages to taking multiple bloodmeals as human hosts are readily available (47–49).

A determining factor of dehydration-induced refeeding is the ability of mosquitoes to locate a resting site with relatively high humidity (this study) and moderate temperature (50). If unable to locate a suitable resting site, dehydration will be rapid under warm and dry conditions, leading to an increased likelihood of multiple bloodmeals. Importantly, as conditions within human dwellings are commonly drier or have less access to free water than in outdoor environments (51), locating a resting site that suppresses dehydration-induced refeeding may be difficult. In fact, our large cage assays held under household conditions (33 ± 4% RH and 23 ± 2°C), confirmed that nearly 50% of *Ae. aegypti* with access to water sources, and 100% of *Ae. aegypti* without access to water sources would take a secondary bloodmeal after feeding to repletion in the previous three days. In more confined spaces (medium cage assays), even *Ae. aegypti* held in considerably more favorable conditions (75 ± 4% RH and 26 ± 2°C) saw similar 50%, and 100% rates of refeeding when held with, and without access to water sources, respectively. Increased feeding in both drier and more confined spaces could explain the high prevalence of multiple bloodmeals when mosquitoes are collected within a household, especially when compared to lower observations of multiple feeding in outdoor biomes (19, 22, 23, 26). Interestingly, ecological factors of a similar type, dry season intensity and human population density, were found to be the primary drivers of human odor preference variation in *Ae. aegypti* mosquitoes (48). It follows then, that if mosquitoes rest in an indoor or outdoor biome, warmer conditions, low humidity, and a lack of water sources will likely lead to substantial refeeding events, yielding two or three bloodmeals during a single gonotrophic cycle. These findings, coupled with intertwined rates of increasing urbanization and predicted mosquito evolution, indicate that mosquitoes will continue to shift towards anthropophilic behaviors (48). Importantly, our observations of multiple bloodmeals improving survival could extend beyond a single gonotrophic cycle, allowing adult mosquitoes to survive for weeks at a time without access to water.

Our findings on compensatory feeding are consistent with previous research that found small, undernourished female *Ae. aegypti* could initially use a bloodmeal for follicle development and then again for ovary maturation, whereas large, properly nourished females could use a single bloodmeal for ovary maturation (20, 52). Furthermore, refeeding may be necessary if the first bloodmeal is interrupted. The flexibility of these mosquitoes to forgo reproduction in exchange for survival or increased nutritional reserves (53) permits them to refeed with only minimal physiological impacts (20, 52, 53). Previous observations and those in this study of multiple feeding events have shown that the process of oogenesis has commenced before a second or third bloodmeal during the same gonotrophic cycle (19, 20). Here, we show little impact on oocyte size or reproductive output in egg number following a second dehydration-induced bloodmeal, so long as mosquitoes can find a location to lay eggs even after 20 days of retaining the eggs. Specific genes have been previously linked with egg retention during drought periods in *Ae. aegypti (38)*, which supports these studies that eggs can be retained under suboptimal conditions until a water source can be located to deposit eggs. Our studies suggest that the second bloodmeal is not directly required for oogenesis; instead, the primary role is to ensure hydration and survival until eggs can be deposited.

Following a bloodmeal, host sensing is suppressed among most mosquito species until eggs are deposited (26, 54–57) but exceptions are known. For example, *Anopheles* mosquitoes have been shown to have increased behavioral attraction to the host following a bloodmeal (26, 54). Suppression of blood feeding is due to a combination of three specific factors - mechanosensation of the ingested blood due to abdominal distension, bloodmeal digestion and processing, and egg development (26, 54). Here, we observe that mosquitoes held under dehydrating conditions resume blood feeding activity and host attraction two-to-three days after an initial bloodmeal. Since this increased blood feeding occurs during oocyte development, the cause is likely associated with improper nutrient-sensing or altered levels of hormones that regulate vitellogenesis and oogenesis processes (see reviews (20, 26)). Another potential mechanism is that factors underlying water-seeking may be altered and increase host detection (58, 59). Interestingly, a return to blood feeding also occurs in mutants with altered humidity detection (Ir8a and Ir93a, (39, 40, 60)), suggesting that our observations are not solely the result of increased water detection and ingestion. Previous studies have shown that dehydrated mosquitoes are remarkably efficient at reequilibrating hemolymph osmolality following a bloodmeal (61), and that dehydration prompts general dysfunction in circulating hemolymph levels through increased osmolarity and altered heart rates (62), which likely results in direct impacts to behavioral and physiological processes. As an example, female mosquitoes detect CO_2_ with a specific class of olfactory receptor neurons (ORNs) designated cpA that express three conserved members of the gustatory receptor (Gr) gene family associated with CO_2_ detection (44, 63). A possibility is that cpA is reactivated to detect changes in CO_2_ during dehydration, allowing mosquitoes to recover sensitivity to this host cue after blood feeding. Additional studies will be required to establish the specific mechanism for how dehydration triggers increased activity and CO_2_ sensing in mosquitoes.

Dry periods have been associated with increased viral transmission (9, 10). This increase was associated with a multitude of factors, which range from altered mosquito-predator interactions associated with temporal water pools (14), altered blood feeding propensity (5), and altered viral dissemination in the mosquito (7, 45, 64). We expand on these observations to show that dry conditions will increase refeeding, which is commonly observed in field-collected mosquitoes (19, 20). Chronic feeding under dry conditions is predicted to substantially increase mosquito survival and elevate blood feeding, prompting predicted increases in vectorial capacity and viral transmission. Although alternative water sources may reduce chronic feeding during dry periods, such as the increased sugar feeding observed in *Ae. albopictus* (36), it is known that standing water sources are limited and the water content of nectar is reduced during drought (5, 14, 37), indicating that increased interactions with a host to blood feed could represent the most reliable means of water acquisition. This could explain why as much as half of field collected mosquitoes had ingested multiple bloodmeals when collected during dry seasons (19, 23).

### Conclusions

Dehydration stress has repeatedly been implicated in water and nutrient depletion, and compensatory mechanisms are known to be utilized to offset the detriments (5, 6, 32). Unfortunately, many of those mechanisms operate through blood feeding, likely resulting in altered disease propagation dynamics within the vector and through host-vector interactions. In addition to postulating the effects on disease transmission, the characterization of refeeding behaviors in dehydrated mosquitoes is essential for determining influences on survival, reproduction, as well as rehydration during and after dehydrating conditions. As climate change is predicted to drive more stark contrasts between wet and dry conditions (16), understanding how these dynamics alter mosquito biology is critical. This study continues to build on how environmental factors, especially drought-like conditions, alter the behavior and physiology of mosquitoes, and ultimately influence disease transmission dynamics.

## Materials and Methods

### Mosquito husbandry

Standard practices were used for rearing and containment of *Ae. aegypti* and *An. stephensi* mosquitoes. Larvae were fed ground fish food (Tetramin) with the addition of yeast extract (Fisher). Adults were maintained in 30 x 30 x 30 cm cages (BugDorm), under a 16h:8h light:dark cycle, with unlimited access to cotton wicks soaked in deionized water and a 10% sucrose solution, all under insectary conditions at approximately 80% relative humidity (RH) and 27°C (vapor pressure deficit (VPD) = 0.71 kPa), unless otherwise specified.

### Mosquito refeeding

Resting conditions between bloodmeals were used to represent conditions that would be common in human dwellings (19, 21, 23). Refeeding was tested under three scenarios - in small cages with single mosquitoes, and in both medium and large cages with ten mosquitoes per cage. For small cages, individual mosquitoes were placed in 17.5 x 17.5 x 17.5 cm cages (BugDorm) held at 33 ± 4% RH and 26 ± 2°C (low humidity; VPD = 2.34 kPa) and two at 75 ± 4% RH and 26 ± 2°C (high humidity; VPD = 0.87 kPa) with access to water. For medium cages, 10 adult mosquitoes (7-14 days old) were placed into four 30 x 30 x 30 cm cages (BugDorm), two with access to DI water and 10% sucrose solution *ad libitum* and two without (+/-), before the cages were placed into separate humidity-controlled 60-quart plastic containers with lids. Four plastic containers were used, two held at low and high humidity. This design yielded four experimental conditions: Two cages of 10 mosquitoes at 33% RH, one with access to water and sucrose solutions *ad libitum* (33+) and one without (33-), and two cages at 75% RH, one with water and sucrose solutions (75+), and one without (75-). Mosquitoes were placed into these experimental containers for 18 hours before being provided the opportunity to blood feed on a live human host for approximately 10 minutes (27-year-old male, leg inside the cage; IRB 2021-0971, University of Cincinnati). A feeding opportunity was presented every 12 hours after the initial feeding for a total of three days. After the 72-hour feeding period, all experimental cages remained within their respective humidified chambers but were supplemented with water, 10% sucrose solution, and oviposition dishes. Mosquitoes were permitted to lay eggs for one week before total egg counts were determined. In addition, in the small cage assays, mosquitoes were dissected, and the ovariole number was assessed to ensure that there were no changes in the number of progeny generated due to multiple feeding events in our experiments.

During the prolonged exposure experiment, mosquitoes were held under dry conditions (75% RH, no access to water) with or without access to water and sugar and allowed to blood feed every 48 hours as survival was assessed. After 20 days, all experimental cages were provided access to water, 10% sucrose solution, and oviposition dishes. The mosquitoes were permitted to lay eggs for one week before total egg counts were determined.

Groups of 10 adult mosquitoes were collected into two 1.8 x 1.8 x 1.8 m cages (BioQuip) kept at 33 ± 4% RH and 23 ± 2°C (VPD = 1.99 kPa), one with access to DI water and 10% sucrose solution *ad libitum* and one without (+/-). Mosquitoes were placed into these experimental cages for 18 hours before being provided the opportunity to bloodfeed on a live human host for approximately 15 minutes (three 27-to 40-year-old male volunteers, leg inside the cage; IRB 2021-0971, University of Cincinnati). A feeding opportunity was presented every 24 hours after the initial feeding for a total of three days. These assays were repeated with the use of mutant mosquito lines to assess refeeding potential, which included the following mutants: ionotropic receptor 8a (39), gustatory receptor 3 - Gr3 (41), Ir93a (40), and odorant receptor co-receptor - Orco (42) to assess host cue detection. The time until each mosquito bite was recorded to determine the time until initial feeding (bites on day 1) and time until refeeding (days 2-4).

### Activity and sleep measurements after blood feeding

The rest-activity rhythms of the mosquitoes were quantified with the aid of a Locomotor Activity Monitor 25 (LAM25) system (TriKinetics Inc., Waltham, MA, USA) and the DAMSystem3 Data Collection Software (TriKinetics) based on methods developed for mosquitoes (65–67). Individuals were blood-fed (27-year-old, male volunteer, leg inside the cage; IRB 2021-0971, University of Cincinnati) before being transferred to glass tubes simulating wet or dry conditions, with sponges soaked in DI water or with dry sponges, respectively. The glass tubes were then positioned horizontally in the LAM25 system, allowing the simultaneous recording of 32 individuals in an “8 x 4” horizontal by vertical matrix. Replicates were intermixed between trials to allow for randomization. The entire set-up was placed in a secluded incubator at 24°C, 70-75% RH, under a 12hr:12hr L/D cycle. After 3-6 hours (coinciding with the start of the night phase), activity level was measured in 1 min bins (the number of times an individual crosses an infrared beam) for three days. Sleep and activity levels were assessed as previously described in mosquitoes (65–67). For comparison with unfed mosquitoes, some individuals were included in the set-up as described above, and data was also retrieved from a previous study (65). Data collected with the DAMSystem3 was processed using the Rethomics platform in R with associated packages, including *behavr*, *ggetho*, *damr,* and *sleepr* (68).

### Uniport assays for blood feeding assessment

A modified uniport assay was developed based on Castillo et al. (69). Briefly, three mosquito cages (30 x 30 x 30 cm, BugDorm) were connected by a 10 cm i.d. acrylic tube (10 cm in length with removable covers to prevent mosquito movement). Mosquitoes were initially confined to the first cage, provided a bloodmeal, and held under conditions that allowed for hydration (30-40% RH with access to multiple water sources) or promoted dehydration (30-40% RH without access to water sources). After two days, mosquitoes were allowed access to all three chambers, and a human host or host mimic (Hemotek) was added to the third chamber. After 5, 10, and 30 minutes, the number of mosquitoes that had successfully started blood feeding was assessed.

### Vectorial capacity, disease transmission modeling analyses, and statistical analyses

As in a previous study (6), vectorial capacity was calculated to determine fold-changes between wet and dry conditions over the 3-day testing period. Egg production was determined from our prolonged survival data, and the overall biting rate per day was determined from our large-cage refeeding data. Disease and population growth modeling was determined based on methods previously developed for mosquito survival and viral transmission in relation to drought stress (5). The full description of the modeling and statistical analyses are provided in the supplement (Figure S4-S13, Tables S1-S5).

## CRediT authorship contribution statement

**Christopher J. Holmes:** Conceptualization, Methodology, Software, Validation, Formal analysis, Investigation, Resources, Data curation, Writing – original draft, Writing – review & editing, Visualization, Supervision, Project administration. **Souvik Chakraborty:** Formal analysis, Investigation, Resources, Data curation, Writing – review & editing. **Oluwaseun Ajayi:** Formal analysis, Investigation, Resources, Data curation, Writing – review & editing. **Melissa Uhran:** Investigation, Data curation, Writing – review & editing. **Ronja Frigard:** Investigation, Data curation, Writing – review & editing. **Crystal Stacey:** Investigation, Data curation, Writing – review & editing. **Emily Susanto:** Investigation, Data curation, Writing – review & editing. **Shyh-Chi Chen:** Writing – review & editing. **Jason L. Rasgon:** Writing – review & editing, Project administration, Funding acquisition. **Matthew DeGennaro:** Writing – review & editing, Project administration, Funding acquisition. **Yanyu Xiao:** Conceptualization, Methodology, Validation, Investigation, Resources, Data curation, Writing – review & editing, Supervision, Project administration, Funding acquisition. **Joshua B. Benoit:** Conceptualization, Methodology, Validation, Investigation, Resources, Data curation, Writing – review & editing, Supervision, Project administration, Funding acquisition.

## Supplemental figure legends

**Figure S1 - Dehydration increases refeeding in mosquitoes.** A. Summary of the studies with mosquitoes held under wet (hydrating) and dry (dehydrating) conditions. B. Medium cages with *Ae. aegypti* (30 x 30 x 30 cm, 10 mosquitoes, N=4), and C. Large cages with *An. stephensi* (1.8 x 1.8 x 1.8 m, N=8, 10 mosquitoes per cage). Dry conditions increased refeeding (total bites – initial bites) for both *Ae. aegypti* and *An. stephensi* (ANOVA with Tukey HSD, and Student’s t-test, P < 0.05).

**Figure S2 - Mosquito mass and ovariole changes following a secondary bloodmeal.** A. Summary of the studies with mosquitoes held under wet (hydrating) and dry (dehydrating) conditions. B. Number of eggs produced per *Aedes aegypti* bite over three days of refeeding under dry and wet conditions, with and without access to water and sucrose solution (30 x 30 x 30 cm, 10 mosquitoes, N=4). C. and D. Mass and ovariole counts for *An. stephensi* measured in small cages (10 x 10 x 10 cm, single mosquitoes, N =15-25). E. and F. Ovariole widths for *An. stephensi* and *Ae. Aegypti,* respectively, were measured in small cages (10 x 10 x 10 cm, single mosquitoes, N =15-25). Eggs per bite and ovariole widths differed in *Ae. aegypti, An. stephensi* ovariole widths increased in size, while ovariole counts and mosquito weights did not show consistent differences over three days of refeeding (ANOVA with Tukey HSD).

**Figure S3 - *Ae. aegypti* bite ratios and time until feeding comparisons for mutant lines.** A. Summary of the studies with mosquitoes held under wet (hydrating) and dry (dehydrating) conditions. B. Mean biting ratios between dry and wet treatments for all mutant lines. C. and D. Combined and separate feeding (day 1) and refeeding times (days 2-4) for all mutant lines, wet and dry conditions. E. Ratios between feeding and refeeding times for wet and dry treatments and all mutant lines. Consistent differences were not apparent in the biting, feeding, nor refeeding data (ANOVA with Tukey HSD and confidence interval comparisons for ratios).

## Supplemental modeling methods

Details for mosquito population growth and viral transmission are provided in the supplement for analyses underlying Figure 4.

