## Supplemental materials for "Multiple blood feeding bouts in mosquitoes allow for prolonged survival and are predicted to increase viral transmission during drought"

### Figure S1

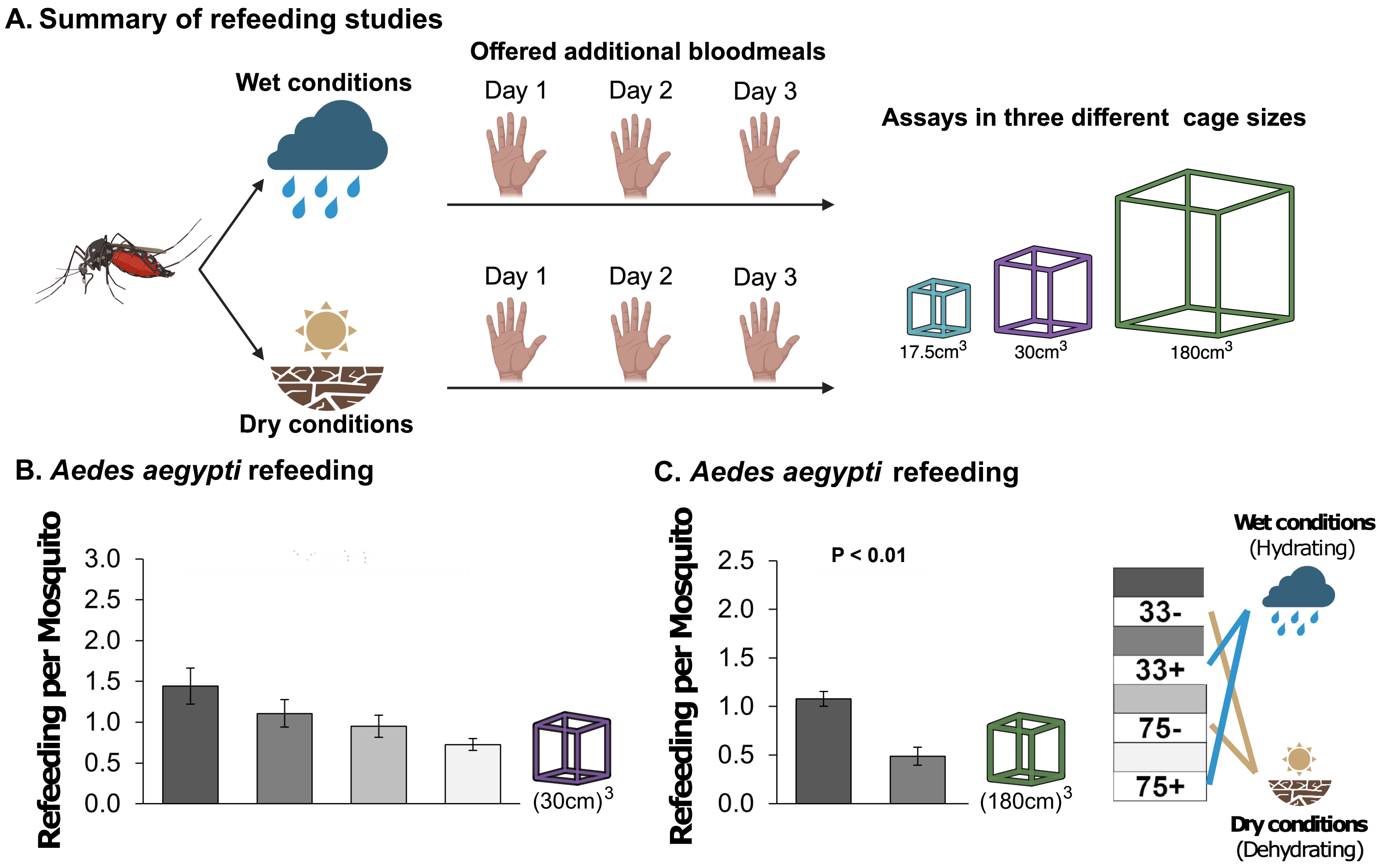
A. Summary of refeeding studies
Offered additional bloodmeals
Wet conditions
Day 1
Day 2
Day 3
Assays in three different cage sizes
Day 1
Day 2
Day 3
17.5cm3
30cm3
180cm3
Dry conditions
B. Aedes aegypti refeeding
Refeeding per Mosquito
3.0
2.5
2.0
1.5
1.0
0.5
0.0
(30cm)3
C. Aedes aegypti refeeding
Refeeding per Mosquito
P < 0.01
2.5
2.0
1.5
1.0
0.5
0.0
(180cm)3
33-
33+
75-
75+
Wet conditions
(Hydrating)
Dry conditions
(Dehydrating)

### Figure S2

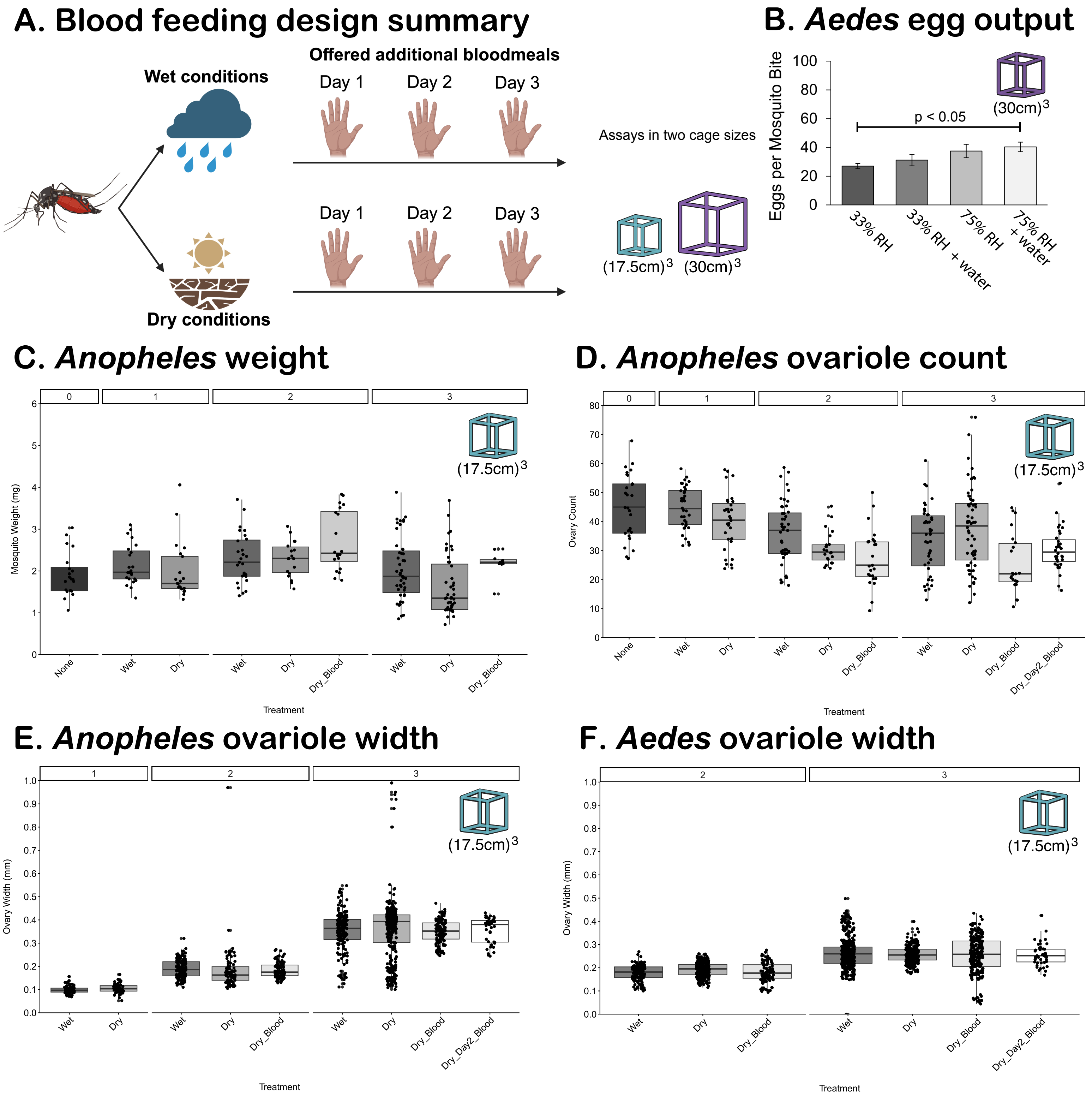
A. Blood feeding design summary
Wet conditions
Offered additional bloodmeals
Day 1
Day 2
Day 3
Dry conditions
Assays in two cage sizes
(17.5cm)3
(30cm)3
B. Aedes egg output
Eggs per Mosquito Bite
p < 0.05
33% RH
33% RH + water
75% RH
75% RH + water
C. Anopheles weight
Mosquito Weight (mg)
None
Wet
Dry
Dry_Blood
Treatment
D. Anopheles ovariole count
Ovary Count
Dry_Day2_Blood
E. Anopheles ovariole width
Ovary Width (mm)
F. Aedes ovariole width

### Figure S3

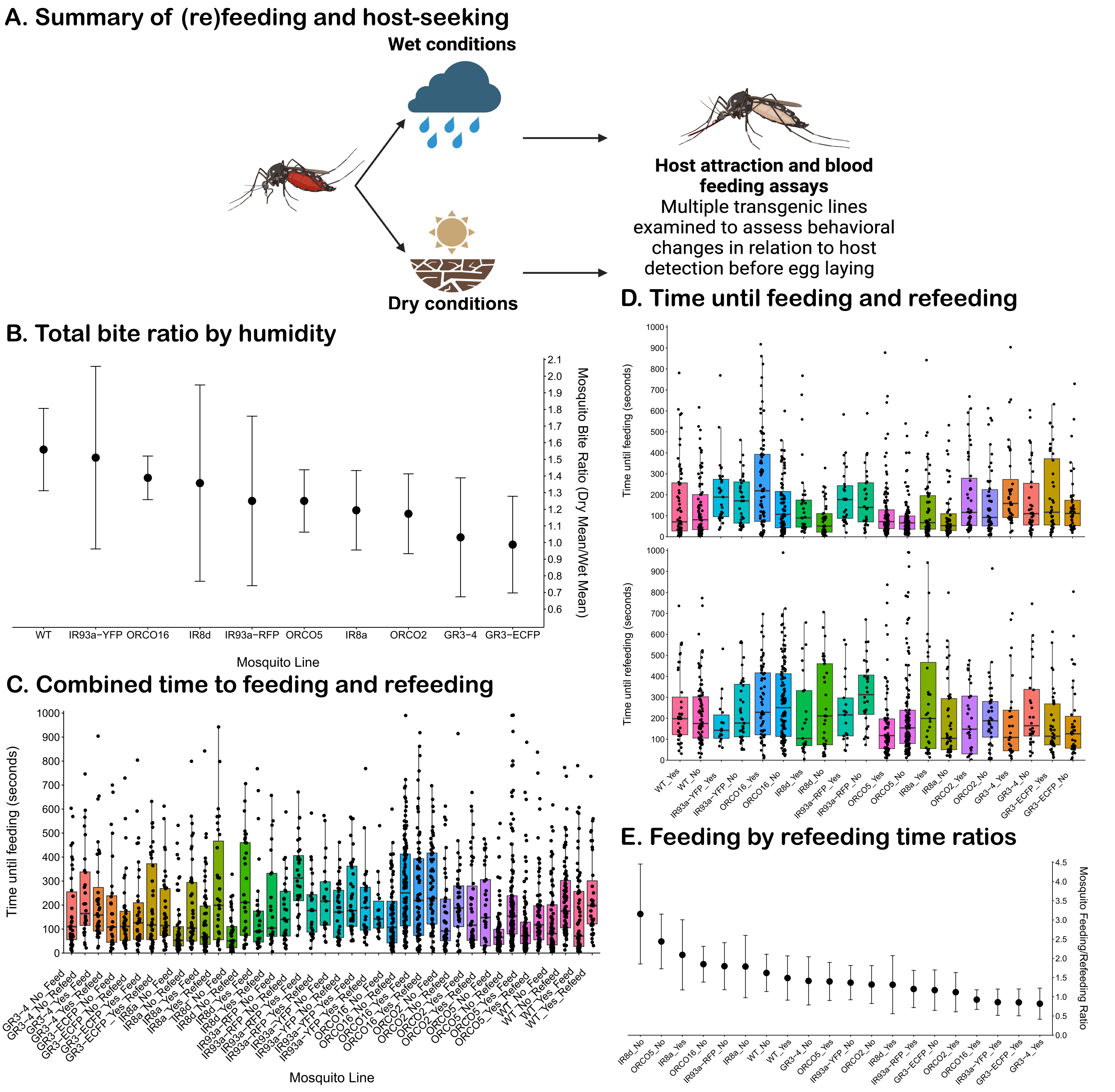
A. Summary of (re)feeding and host-seeking
Wet conditions
Dry conditions
Host attraction and blood feeding assays
Multiple transgenic lines examined to assess behavioral changes in relation to host detection before egg laying
B. Total bite ratio by humidity
Mosquito Bite Ratio (Dry Mean/Wet Mean)
Mosquito Line
WT
IR93a−YFP
ORCO16
IR8d
IR93a−RFP
ORCO5
IR8a
ORCO2
GR3−4
GR3−ECFP
C. Combined time to feeding and refeeding
Time until feeding (seconds)
Mosquito Line
D. Time until feeding and refeeding
Time until feeding (seconds)
Time until refeeding (seconds)
E. Feeding by refeeding time ratios
Mosquito Feeding/Refeeding Ratio

### Supplementary methods

Based on varying environmental pressures encountered during different developmental stages of mosquitoes, we utilized an age-structured model to depict the population dynamics of mosquitoes under fluctuating environmental conditions. We denote fives stages for mosquito development: E for eggs. L for larva (and pupa), $A_1$ for adult mosquitoes experiencing dehydration stress, $A_2$ for adult mosquitoes under dehydration stress with access to bloodmeals, and $A_3$ for adult mosquitoes without dehydration stress and with water access. These adult mosquito statuses, each characterized by distinct survival rates and capable of transitioning between one another, mimic three environmental scenarios: dry seasons in sparsely populated regions, dry seasons in densely populated re-gions, and wet seasons. Following bloodmeals, adult mosquitoes with water access initiate their ovulation period and lay eggs at a rate of $bB(\cdot)$, where $B(\cdot)$ represents the daily birth function determined by time and the number of adult mosquitoes ready for egg laying. Eggs progress to the larval and pupal stages at a rate of $a_1$, then further mature into adult mosquitoes facing the three aforementioned environmental conditions at a rate of $a_2$. Despite the potential for eggs and larvae (pupae) of certain mosquito species to perish quickly under dehydration stress due to their reliance on water, some adult mosquitoes can sustain themselves through bloodmeals (this study). After dehydration stress, female mosquitoes typically delay egg laying for approximately one day to adapt to environmental changes. In Figure S4, the flowchart illustrates the schematic representation of our model depicting the population dynamics of mosquitoes.

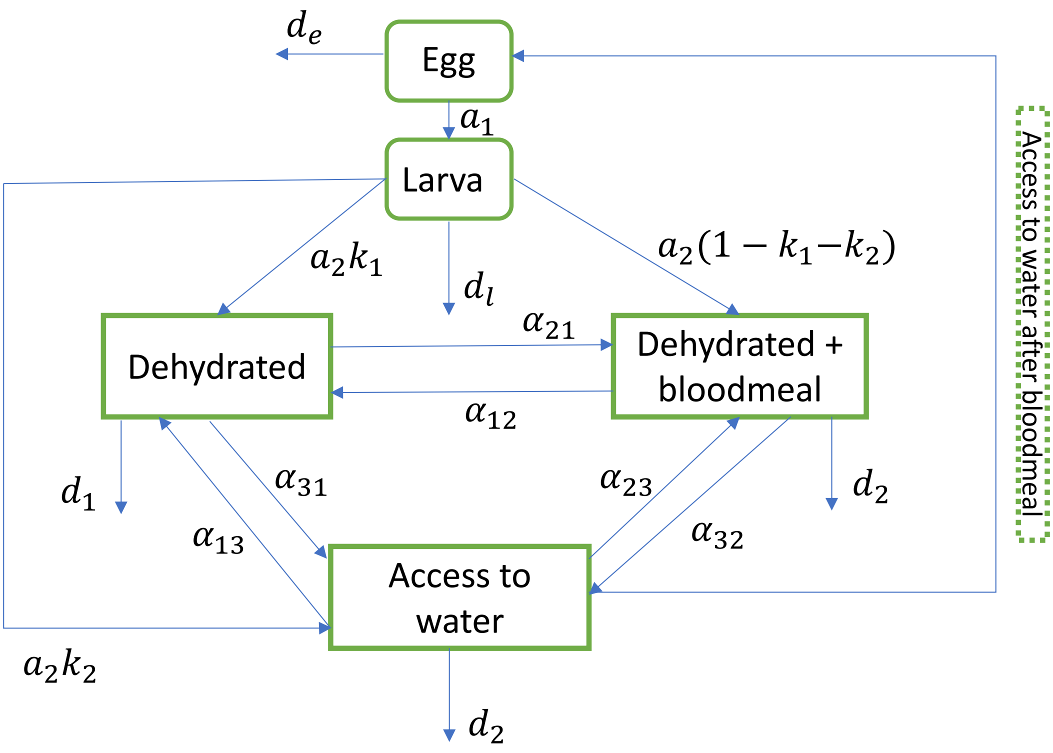


Figure S4: The flowchart for the life cycle of mosquito under environmental pressures.

Based on the flowchart in Figure S4, we developed the model (1) to explore mosquito population dynamics under environmental pressures. The parameters in the model are listed in Table S1 and Table S2.

Table S1: Parameters for mosquito [2]

| Parameter | Descriptions | Range |
|---|---|---|
| $b$ | Probability of egg hatching | See Table S2 |
| $B(\cdot)$ | Birth function | See description in the text |
| $e$ | Average number of eggs laid by a female mosquito | $100[50-200]$ per oviposition period |
| $d_c$ | Death rate of mosquitoes eggs due to intra-species competition | 0.0000001 per day per mosquito |
| $d_I$ | Death rate for larva/pupa | See Table S2 |
| $a_1$ | Developmental rate from eggs to larva/pupa | See Table S2 |
| $a_2$ | Developmental rate from larva/pupa to matured mosquitoes | See Table S2 |
| $k_1$ | Proportion of recently matured mosquitoes in dehydrated status | See Table S2 |
| $k_2$ | Proportion of recently matured mosquitoes with water ac-cess | See Table S2 |
| $\alpha_{ij}$ | Transition rate from status $j$ to $i$, where $i, j$ represents one of the statuses for matured mosquitoes | See Table S2 |
| $d_1$ | Death rate for matured mosquito in dehydrated status | See Table S2 |
| $d_2$ | Death rate for matured mosquitoes in dehydrated status with access to bloodmeals | See Table S2 |
| $d_3$ | Death rate for matured mosquitoes with access to water | See Table S2 |

$$
\begin{aligned}
E' &= bB(\alpha_{32}A_2) - (d_e + a_1 + d_c E)E, \\
L' &= a_1 E - (d_l + a_2)\,L, \\
A_1' &= k_1 a_2 L + \alpha_{12}A_2 + \alpha_{13}A_3 - (d_1 + \alpha_{21} + \alpha_{31})A_1, \\
A_2' &= (1 - k_1 - k_2)a_2 L + \alpha_{21}A_1 + \alpha_{23}A_3 - (d_2 + \alpha_{12} + \alpha_{32})A_2, \\
A_3' &= k_2 a_2 L + \alpha_{31}A_1 + \alpha_{32}A_2 - (d_3 + \alpha_{13} + \alpha_{23})A_3.
\end{aligned} \tag{1}
$$

Here, we have $\alpha_{ij}$ and $d_i$, where $i, j = 1, 2, 3$ representing the transition rate of adult mosquitoes from status $j$ to status $i$ and death rate of mosquitoes, which are environmental dependent, i.e., during the dry period, mosquitoes would experience high mortality rate due to lack of water and switch to a status that relies on bloodmeals to survive longer. We also have the birth function for average number of egg laid by a mosquito per day as

$$
B(\alpha_{32}A_2) = \frac{e\alpha_{32}A_2}{14},
$$

where $e$ is the average number of eggs laid by a female mosquito during each oviposition period when allowed access to water (around 14 days). We have the initial condition for mosquito populations as $E(0) = 100$, $L(0) = 100$, $A_1(0) = 1000$, $A_2(0) = 1000$, $A_3(0) = 1000$.

In this study, we investigate the population dynamics of two mosquito species, *Aedes aegypti* and *Culex pipiens*, under environmental pressure. *Culex* species shows greater susceptibility to dehydration pressure compared to *Aedes* species across various environmental conditions [7–9]. We have documented distinct survival rates for each developmental stage of both species, as outlined in Table S2 under different environmental circumstances.

During alternating dry and wet periods, the population dynamics of both mosquito species fluctuate in tandem with the changing seasons, as depicted in Figures S5. Each species experiences a cycle of 7 days of dehydration followed by 21 days of a wet period. Over time, both populations gradually decrease, with the *Culex* population eventually dying out and the *Aedes* population surviving at very low levels, as illustrated in Figure S5 (a). However, when they can obtain water from bloodmeals by biting humans or animals, both species have a chance of survival, potentially leading to a significant increase in the Aedes population, as shown in Figure S5 (b). Notably, the ratio between the duration of the dehydrated period and that of the wet period significantly impacts population sustainability, as indicated in Figure S6. After 10 repeated cycles of alternating dry and wet periods, the *Culex* population faces extinction if bloodmeals are not substituted when this ratio falls below 2.5, as demonstrated in Figure S6 (c). However, when the length of the wet period, representing the recovery time for the populations, is at least double that of the dry period, both the *Culex* population with blood meal substitution and the *Aedes*.

Table S2: Environmental dependent variables [1, 2, 5, 6, 10]

| Category | Dry season | | | Wet season | | | Dry season with bloodmeals as substitution | | |
|---|---|---|---|---|---|---|---|---|---|
| Transition rate | - | 0.8 | 0.8 | - | 0.1 | 0.3 | - | 0.1 | 0.1 |
| | 0.1 | - | 0.1 | 0.8 | - | 0.6 | 0.2 | - | 0.3 |
| | 0.1 | 0.1 | - | 0.1 | 0.1 | - | 0.8 | 0.4 | - |
| *Culex* | $d_1 = 1$ | | | $d_2 = 5/30$ | | | $d_3 = 1/30$ | | |
| | $b = 0$ | | | $b = 1$ | | | $b = 0$ | | |
| | $d_e = 1/4$ | | | $d_e = 1/120$ | | | $d_e = 1/120$ | | |
| | $d_l = 1$ | | | $d_l = 0.0163$ | | | $d_l = 0.0163$ | | |
| | $a_1 = 0$ | | | $a_1 = 1/7$ | | | $a_1 = 0$ | | |
| | $a_2 = 0.05$ | | | $a_1 = 1/3$ | | | $a_1 = 0.05$ | | |
| | $k_1 = 1$ | | | $k_1 = 0.2$ | | | $k_1 = 0.4$ | | |
| | $k_2 = 0$ | | | $k_2 = 0.6$ | | | $k_2 = 0.2$ | | |
| | $\alpha_{HM} = 0.4$ | | | $\alpha_{HM} = 0.2$ | | | $\alpha_{HM} = 0.2$ | | |
| | $\alpha_{BM} = 0.4$ | | | $\alpha_{BM} = 0.2$ | | | $\alpha_{BM} = 0.2$ | | |
| | $\epsilon_M = 1/8$ | | | $\epsilon_M = 1/10$ | | | $\epsilon_M = 1/8$ | | |
| *Aedes* | $d_1 = 1/1.5$ | | | $d_2 = 5/30$ | | | $d_3 = 1/30$ | | |
| | $b = 0$ | | | $b = 1$ | | | $b = 0$ | | |
| | $d_e = 1/45$ | | | $d_e = 1/120$ | | | $d_e = 1/120$ | | |
| | $d_l = 1$ | | | $d_l = 0.0163$ | | | $d_l = 0.0163$ | | |
| | $a_1 = 0$ | | | $a_1 = 1/7$ | | | $a_1 = 0$ | | |
| | $a_2 = 0.05$ | | | $a_1 = 1/3$ | | | $a_1 = 0.05$ | | |
| | $\alpha_{HM} = 0.4$ | | | $\alpha_{HM} = 0.2$ | | | $\alpha_{HM} = 0.2$ | | |
| | $\epsilon_M = 1/8$ | | | $\epsilon_M = 1/10$ | | | $\epsilon_M = 1/8$ | | |

population can survive short-term periodic droughts. In such scenarios, the *Aedes* population may grow rapidly with blood meal substitution (Figure S6 (a)).

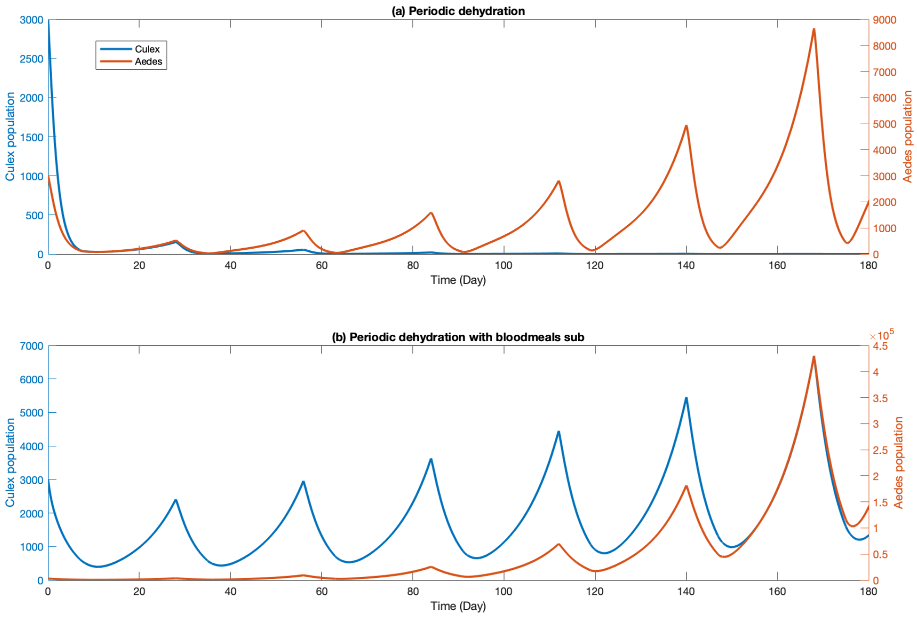


Figure S5: Mosquito population dynamics during environmental pressures. (a) Populations experience 7 days of dehydration and followed 21 days of wet period, then alternating; (b) Same setting as (a) but mosquitoes could get bloodmeals.

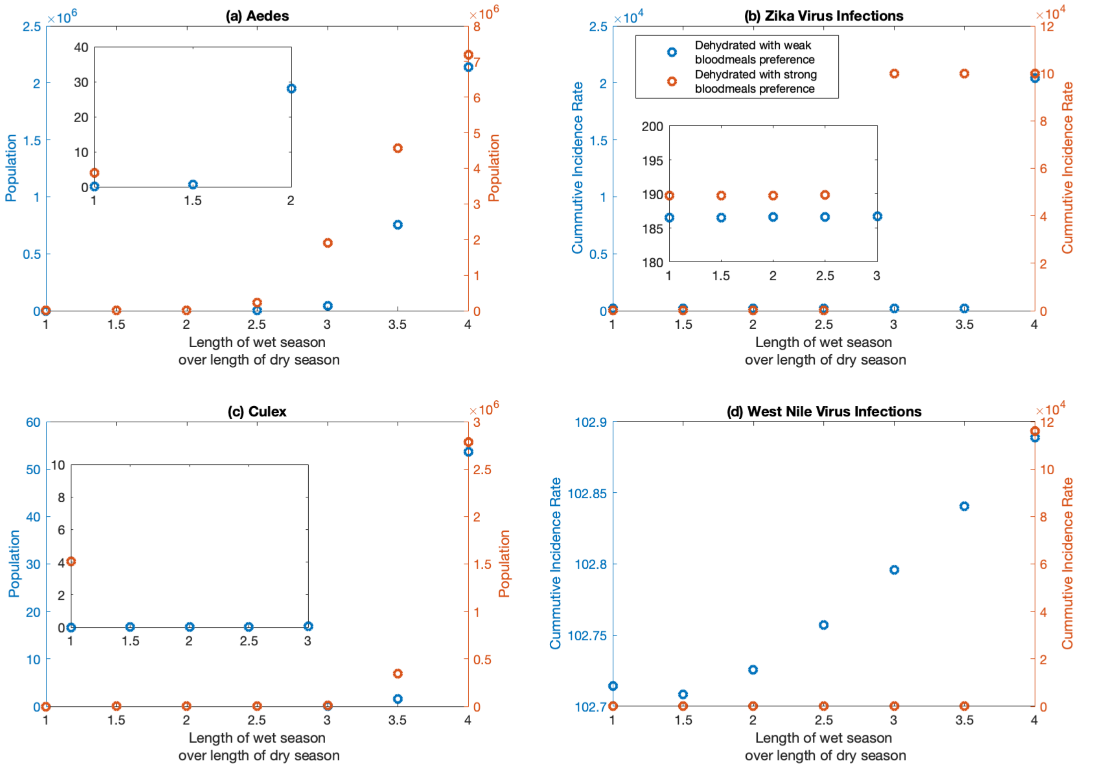


Figure S6: Mosquito population size during environmental pressures and related disease scale after 10 repeated cycles alternating between 7 days of dry period and various ratio of two periods. (a) Population of *Aedes* mosquito;(b) Cumulative number of human infections for Zika virus per 100,000 population at the end of the 10 repeated cycles; (c) Population of *Culex* mosquito; (d) Cumulative number of human infections for West Nile virus per 100,000 population at the end of the 10 repeated cycles.

After examining the population dynamics of mosquitoes under environmental pressure, we further developed an eco-epidemic model to depict the disease dynamics of West Nile Virus and Zika virus infections, alongside the population dynamics of *Culex* and *Aedes* mosquitoes, which are vectors for these two diseases, respectively. For the transmission of West Nile virus, we considered the pathogen transmission among mosquito group $A_2$, humans, and birds.

Building upon previous model frameworks [2, 3, 10, 11], we formulated an age-structured mosquito population model nested within the disease model for West Nile Virus transmission. This model incorporates three human (mosquitoes, birds) compartments: $S_H$ $(S_M, S_B)$ for susceptible humans (mosquitoes, birds), $E_H$ $(E_M, E_B)$ for exposed humans (mosquitoes, birds), and $I_H$ $(I_M, I_B)$ for infectious humans (mosquitoes, birds). We assume that the three compartments of mosquitoes consist of mosquitoes in group $A_2$ in the mosquito model (1), with a death rate for mosquitoes denoted as $d_2$. The model is presented in (2).

Table S3: Parameters for West Nile Virus transmission [1, 2, 10, 11]

| Parameter | Descriptions | Value |
|---|---|---|
| $\eta_h$ | Birth rate for human | 0.3 (assumed) |
| $\alpha_{HM}$ | Probability of pathogen transmission from mosquitoes to human | See Table S2 |
| $\beta_{HM}$ | Mosquito biting rate on human (number of bites per mosquito per day) | 0.262 |
| $d_H$ | Death rate for human | 1/65/365 per day (assumed) |
| $\gamma_H$ | Recover rate | 1/14 per day |
| $d_{HH}$ | Disease related mortality rate of human | 0.05/365 per day |
| $\epsilon_H$ | Disease progressing rate from exposed stage to infectious stage for human | 1/14 per day |
| $\alpha_{MH}$ | Probability of pathogen transmission from human to mosquitoes | 0.8 |
| $\epsilon_M$ | Disease progressing rate from exposed stage to infectious stage for mosquitoes | See Table S2 |
| $d_2$ | Death rate for mosquito looking for bloodmeals | See Table S2 |
| $\eta_B$ | Migration rate for birds | 12 per day (assumed) |
| $b_B$ | Birth rate for birds | 0.01 per day |
| $d_B$ | Death rate for birds | 0.001 per day |
| $K_B$ | Carrying capacity for birds | 1200 (assumed) |
| $\alpha_{BM}$ | Probability of pathogen transmission from mosquitoes to birds | See Table S2 |
| $\beta_{BM}$ | Mosquito biting rate on bird (number of bites per mosquito per day) | 0.262 |
| $\gamma_B$ | Recover rate for birds | 0.001 per day |
| $d_{BB}$ | Disease related mortality rate of birds | 0.143 per day |
| $\epsilon_B$ | Disease progressing rate from exposed stage to infectious stage for birds | 1/10 per day |

$$
\begin{aligned}
S'_H =& \eta_h - \alpha_{HM}\beta_{HM}\frac{S_H}{P_H}I_M - d_H S_H + \gamma_H I_H, \\
E'_H =& \alpha_{HM}\beta_{HM}\frac{S_H}{P_H}I_M - (d_H + d_{HH} + \epsilon_H)E_H, \\
I'_H =& \epsilon_H E_H - (d_H + d_{HH} + \gamma_H)I_H, \\
E'_M =& \alpha_{MH}\beta_{HM}(A_2 - E_M - I_M)\frac{I_H}{P_H} - (d_2 + \epsilon_M)E_M, \\
I'_M =& \epsilon_M E_M - d_2 I_M, \\
S'_B =& \eta_B + b_B(S_B + E_B + I_B) - (b_B - d_B)\frac{(S_B + E_B + I_B)^2}{K_B} - \alpha_{BM}\beta_{BM}S_B\frac{I_M}{P_B} - d_B S_B + \gamma_B I_B, \\
E'_B =& \alpha_{BM}\beta_{BM}I_M\frac{S_B}{P_B} - (d_B + d_{BB} + \epsilon_B)E_H; \\
I'_B =& \epsilon_B E_B - (d_B + d_{BB} + \gamma_B)I_B;
\end{aligned}
\tag{2}
$$

In this context, we have considered constant recruitment for humans and logistic growth functionality for birds. All parameters are detailed in Table S3.

To model Zika virus transmission by *Aedes* mosquitoes, we have considered both human-to-vector-human and human-to-human pathogen transmission pathways. We define the following human subgroups: susceptible ($S_H$), exposed ($E_H$), asymptomatically infected ($A_H$), symptomatically infected ($I_{H1}$), convalescent ($I_{H2}$), recovered ($R_H$), and the total population ($P_H = S_H + E_H + A_H + I_{H1} + I_{H2} + R_H$). The embedded mosquito subgroups follow similar settings as in our West Nile virus model. The complete model for Zika transmission is described by the equations in (3), and the related parameters are listed in Table S4.

Table S4: Parameters for Zika Virus transmission [2–6]

| Parameter | Descriptions | Value |
|---|---|---|
| $\eta_h$ | Birth rate for human | 0.3 (assumed) |
| $\alpha_{HM}$ | Probability of pathogen transmission from mosquitoes to human | See Table S2 |
| $\beta_{HM}$ | Mosquito biting rate (number of bites per mosquito per day) | 0.5 |
| $\alpha_{HH}$ | Pathogen transmission probability between human to human | 0.05 |
| $\kappa$ | Reduced infectivity factor for exposed stage to human | 0.6 |
| $\tau$ | Reduced infectivity factor for asymptomatic infections to human | 0.3 |
| $d_H$ | Death rate for human | 1/65/365 per day (assumed) |
| $\theta$ | Proportion of infections developing to symptomatic infections | 0.18 |
| $\epsilon_H$ | Disease progressing rate from exposed stage to infectious stage for human | 1/5 per day |
| $\gamma_{H1}$ | Recover rate to convalescent stage | 1/7 per day |
| $\gamma_{H2}$ | Recover rate from convalescent stage to recovered stage | 1/5 per day |
| $\gamma_{H3}$ | Recover rate from asymptomatic infections to recovered stage | 1/20 per day |
| $\alpha_{MH}$ | Probability of pathogen transmission from human to mosquitoes | 0.5 |
| $\epsilon_M$ | Disease progressing rate from exposed stage to infectious stage for mosquitoes | See Table S2 |
| $\eta$ | Reduced infectivity factor from $E_H, A_H$ to mosquito | 0.1 |
| $d_2$ | Death rate for mosquito looking for bloodmeals | See Table S2 |

$$
\begin{aligned}
S'_H &= \eta_h - \alpha_{HM}\beta_{HM}I_M\frac{S_H}{P_H} - \alpha_{HH}(\kappa E_H + I_{H1} + \tau I_{H2})\frac{S_H}{P_H} - d_H S_H, \\
E'_H &= \theta(\alpha_{HM}\beta_{HM}I_M\frac{S_H}{P_H} + \alpha_{HH}(\kappa E_H + I_{H1} + \tau I_{H2})\frac{S_H}{P_H} - (d_H + \epsilon_H)E_H, \\
I'_{H1} &= \epsilon_H E_H - (d_H + \gamma_{H1})I_{H1}, \\
I'_{H2} &= \gamma_{H1}I_{H1} - \gamma_{H2}I_{H2}, \\
A'_H &= (1-\theta)(\alpha_{HM}\beta_{HM}I_M\frac{S_H}{P_H} + \alpha_{HH}(\kappa E_H + I_{H1} + \tau I_{H2})\frac{S_H}{P_H} - \gamma_{H3}A_H, \\
R'_H &= \gamma_{H2}I_{H2} + \gamma_{H3}A_H, \\
E'_M &= \alpha_{MH}\beta_{MH}(A_2 - E_M - I_M)\frac{I_{H1} + \eta E_H + \eta A_H}{P_H} - (\alpha_{22} + \epsilon_M)E_M, \\
I'_M &= \epsilon_M E_M - \alpha_{22}I_M
\end{aligned}
\tag{3}
$$

According to the definitions and formulations of the basic reproduction number in [3, 12–14], we have calculated that the $R_0$ values for Zika and West Nile viruses in our presented model settings are always less than 1, except in cases where the ratio of the length of the dry period to that of the wet period is greater than 2.5 for West Nile virus transmission. This indicates that the diseases will eventually die out in the population.

We observed that for the mosquito population without blood meal substitution during periodic dry periods, the cumulative incidence rate is low, and the scale of Zika outbreak remains low when the ratio of the length of the dry period to that of the wet period is no greater than 3, as shown in Figures S6 (b) and S7. However, mosquito populations with blood meal substitution could rapidly escalate the scale of Zika outbreaks to their maximum level after the ratio reaches 2.5, as shown in Figures S6 (b) and 8.

Regarding the *Culex* species, as illustrated in Figures S6 (c) and S7, the mosquito population will either die out survive at very low levels, resulting in a very limited contribution to West Nile virus transmission. If the *Culex* species receives support from bloodmeals during dry periods, they can contribute to boosting the scale of disease outbreaks when the ratio of the two periods is larger than 3, as demonstrated in Figures S6 (b) and S7.

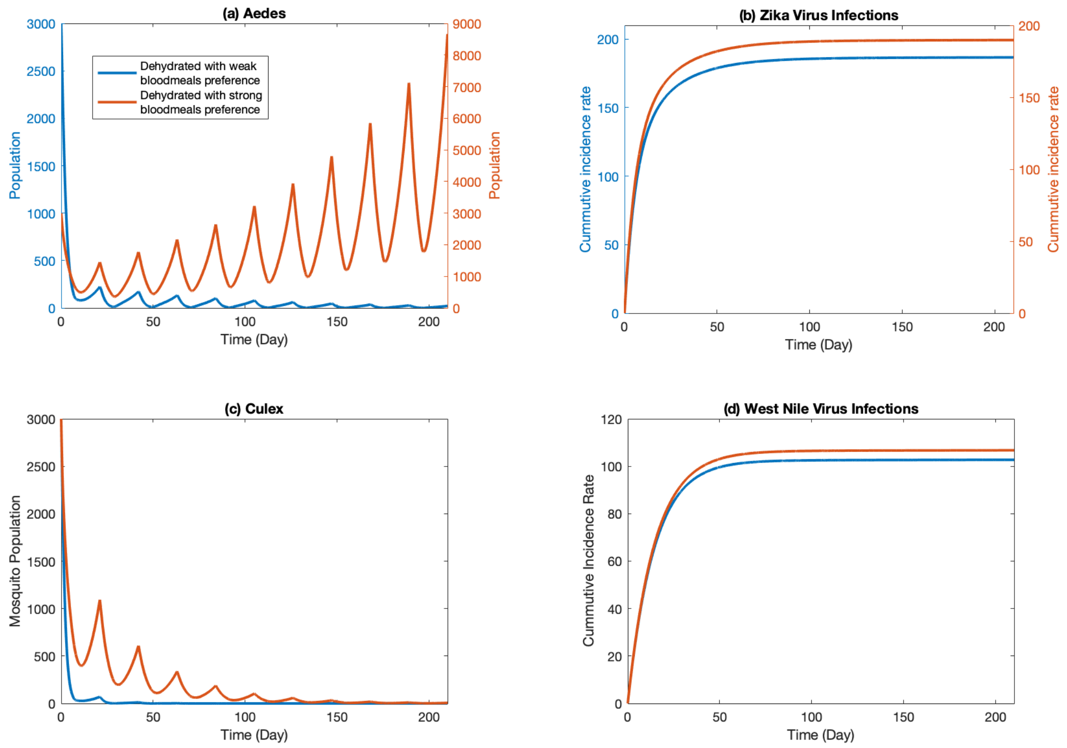


Figure S7: Mosquito population dynamics and related disease scale for the environmental setting of repeated cycles alternating between 7 days of dry period followed by 14 days of wet period. (a) Population of*Aedes* mosquito; (b) Cumulative number of human infections for Zika virus per 100,000 population; (c) Population of *Culex* mosquito; (d) Cumulative number of human infections for West Nile virus per 100,000 population.

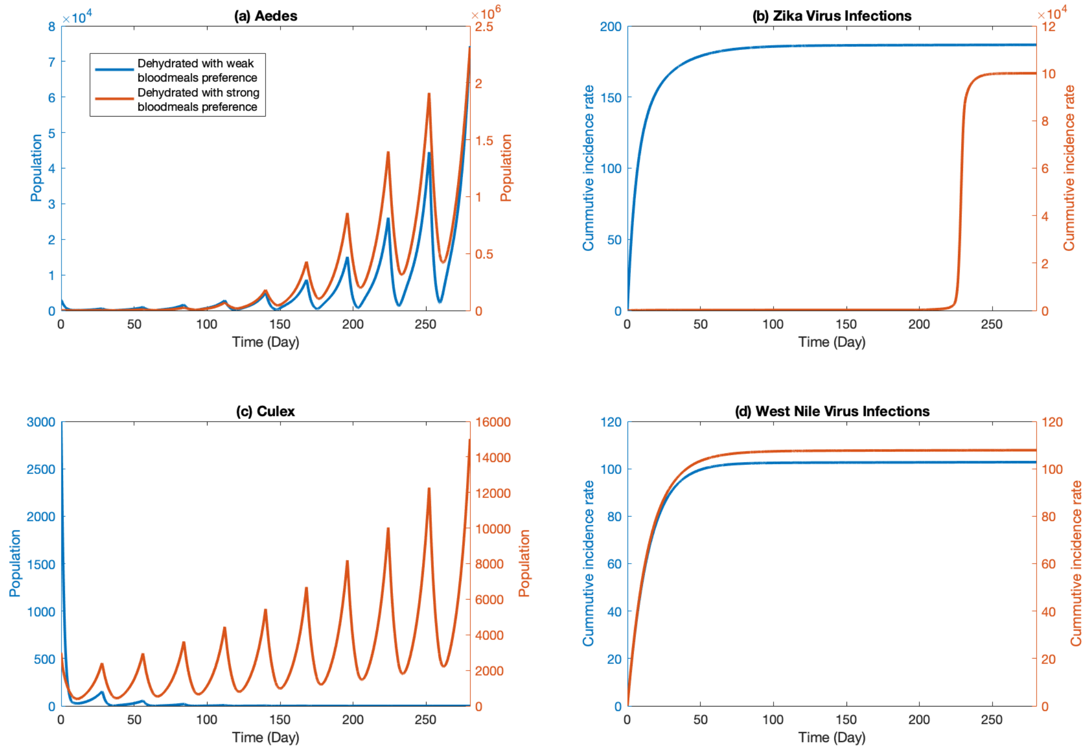


Figure S8: Mosquito population dynamics and related disease dynamics for the environmental setting of repeated cycles alternating between 7 days of dry period followed by 21 days of wet period. (a) Population of *Aedes* mosquito; (b) Cumulative number of human infections for Zika virus per 100,000 population; (c) Population of *Culex* mosquito; (d) Cumulative number of human infections for West Nile virus per 100,000 population.

We further examined scenarios where the length of the dry period is 14 days. We observed that the threshold ratios of the length of the dry and wet periods, which sustain the population, decrease for both *Culex* with the substitution of bloodmeals and *Aedes* with or without bloodmeals due to the longer recovery time in the wet period (see Figure S9 (a) and (c)). Consequently, the cumulative incidence rates for Zika virus infection quickly reach saturated levels (see Figure S9 (b)), and those for West Nile virus infection increase rapidly (Figure S9 (d)). However, the cumulative incidence rate for West Nile virus transmitted by *Culex* without bloodmeals is not sensitive to the ratio of the two periods (blue dots in Figure S9 (d)).

We further explored the population and disease dynamics through time series solutions shown in Figures S10 and S11. It is evident that the amplitude of the mosquito population increases with the increase of the ratio of the two periods. Moreover, when the ratio is large (i.e., greater than 3), the basic reproduction number for West Nile virus transmission exceeds 1 due to the significant increment of the *Culex* population relying on bloodmeals during the dry period (Figure S11 (d)). The scenario with the ratio being 2 is presented in Figure S12.

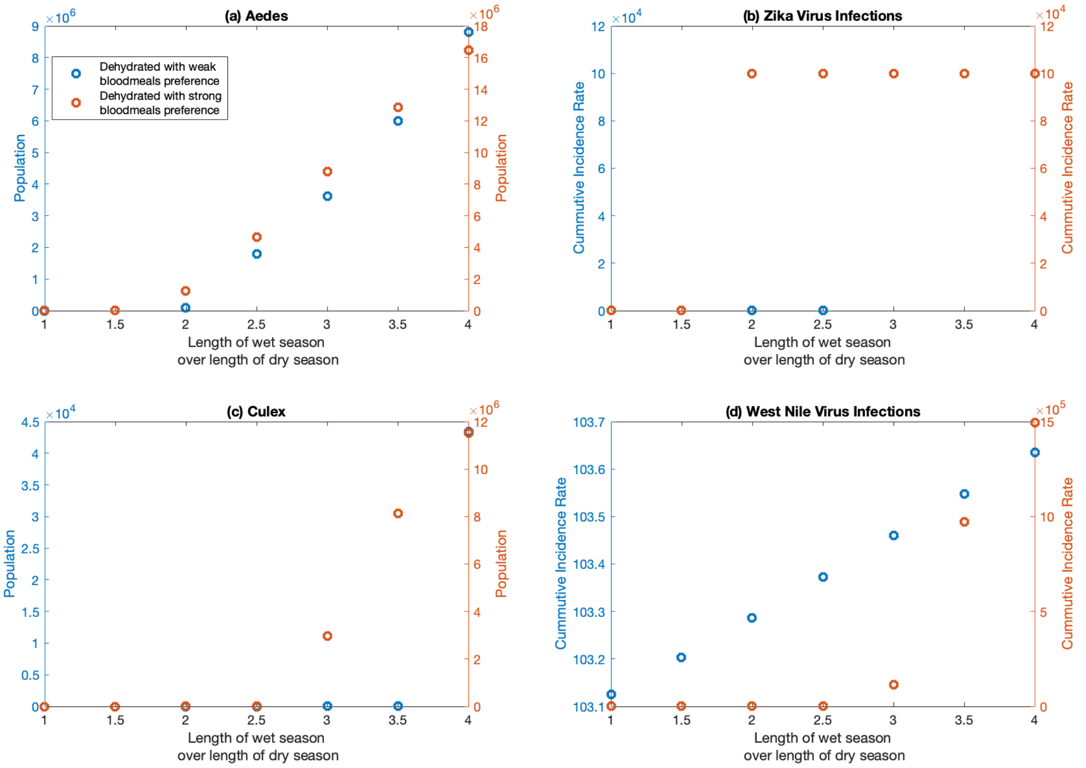


Figure S9: Mosquito population size during environmental pressures and related disease dynamics after 10 repeated cycles alternating between 14 days of dry period and various ratio. (a) Population of *Aedes* mosquito; (b) Cumula-tive number of human infections for Zika virus per 100,000 population at the end of the 10 repeated cycles; (c) Population of *Culex* mosquito; (d) Cumulative number of human infections for West Nile virus per 100,000 population at the end of the 10 repeated cycles.

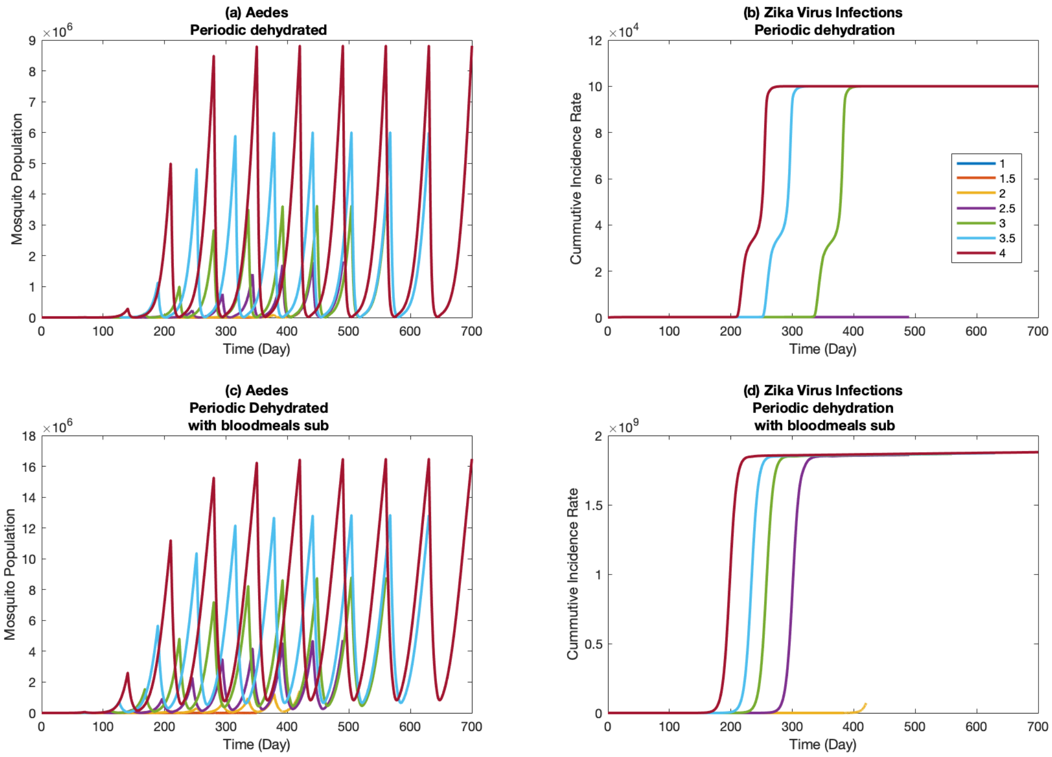


Figure S10: *Aedes* mosquito population dynamics and Zika disease dynamics for the environmental setting of 14 days of dry period and various ratio of two periods. (a) Population of *Aedes* mosquito; (b) Cumulative number of human infections for Zika virus per 100,000 population by mosquito population presented in (a); (c) Population of *Aedes* mosquito with bloodmeals; (d) Cumulative number of human infections for Zika virus per 100,000 population by mosquito population presented in (c).

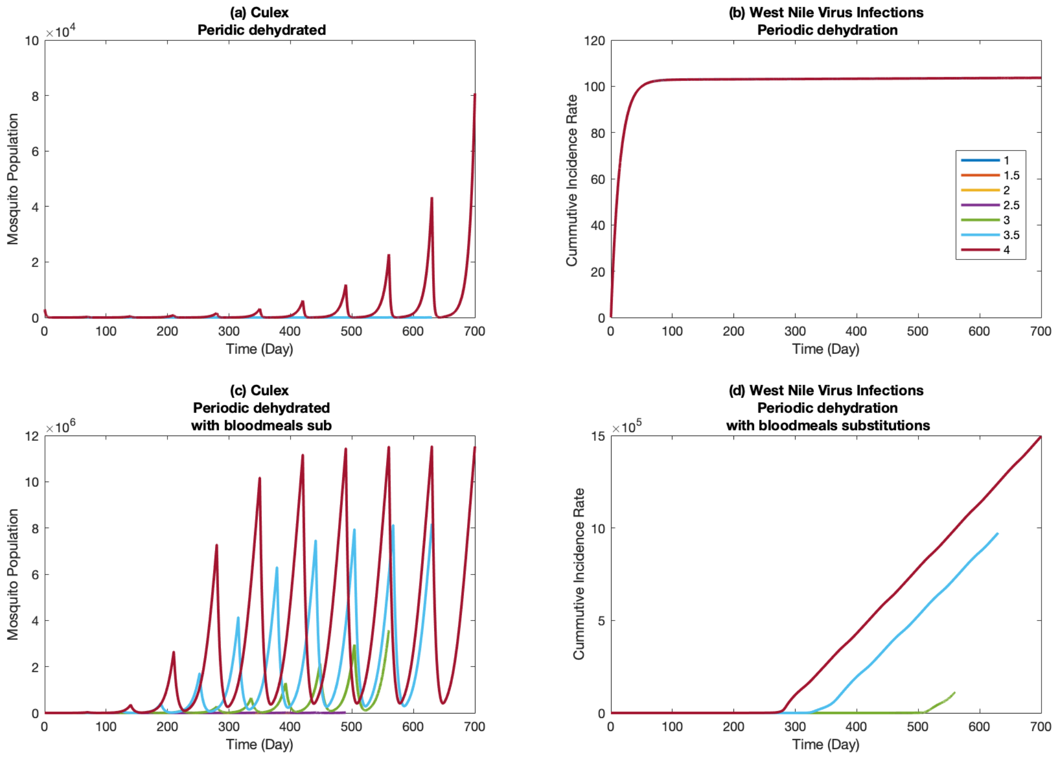


Figure S11: *Culex* mosquito population dynamics and West Nile virus disease dynamics for the environmental setting of 14 days of dry period and various ratio of two periods. (a) Population of *Culex* mosquito; (b) Cumulative number of human infections for West Nile virus per 100,000 population by mosquito population presented in (a); (c) Population of *Culex* mosquito with bloodmeals; (d) Cumulative number of human infections for West Nile virus per 100,000 population by mosquito population presented in (c).

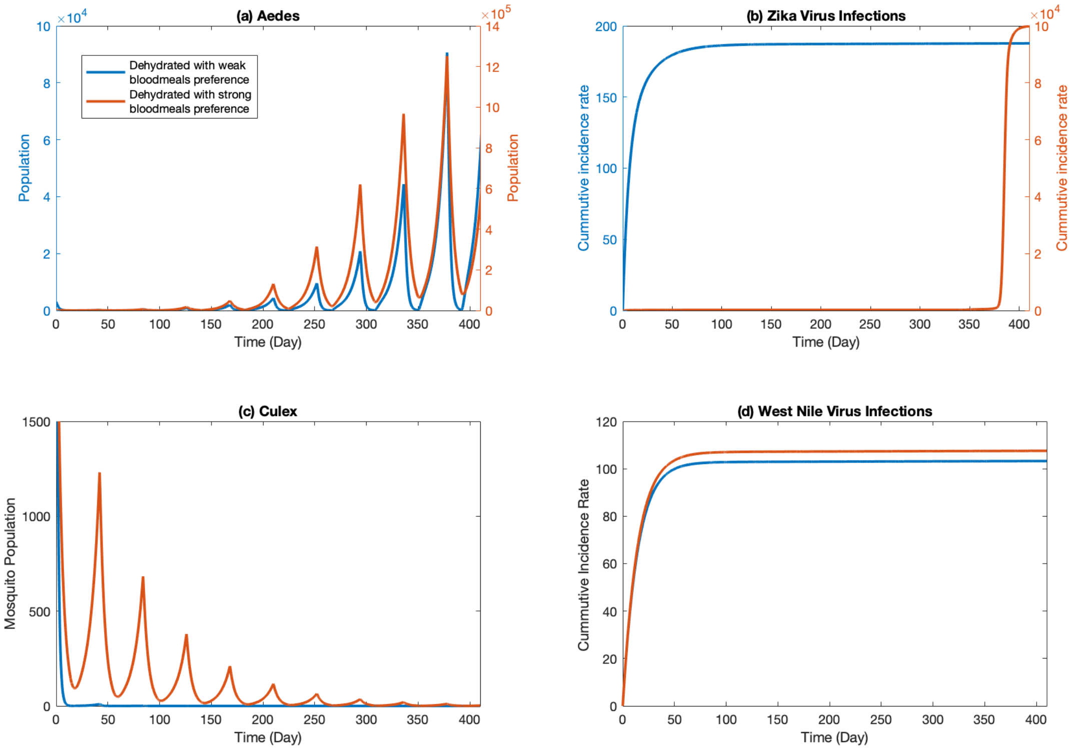


Figure S12: Mosquito population dynamics and related disease dynamics for the environmental setting of repeated cycles alternating between 14 days of dry period followed by 28 days of wet period. (a) Population of *Aedes* mosquito; (b) Cumulative number of human infections for Zika virus per 100,000 population; (c) Population of *Culex* mosquito; (d) Cumulative number of human infections for West Nile virus per 100,000 population.

In addition, we would explore two scenarios: fast and slow physiological responses of mosquitoes towards environmental pressure, where dehydrated mosquitoes exhibit varying degrees of preference for bloodmeals. In this case, we have the transition rates $\alpha_{ij}$ listed in Table S5. We observe that the threshold ratio of the two periods for the survival of mosquito populations increases, and the population sizes are smaller for the case with weaker preference compared to the stronger preference case (see Figures S13 (a)(c) and S14 (a)(c)). However, the related disease scales are more or less the same for the two scenarios (see Figures S13 (b)(d) and S14 (b)(d)).

Table S5: Transition rates for slow reaction of physiology change of mosquito.

| Category | Dry season | | | Wet season | | | Dry season with bloodmeals as substitution | | |
|---|---|---|---|---|---|---|---|---|---|
| Transition rate | - | 0.8 | 0.8 | - | 0.1 | 0.7 | - | 0.1 | 0.1 |
| | 0.1 | - | 0.1 | 0.4 | - | 0.3 | 0.2 | - | 0.3 |
| | 0.1 | 0.1 | - | 0.1 | 0.1 | - | 0.8 | 0.4 | - |

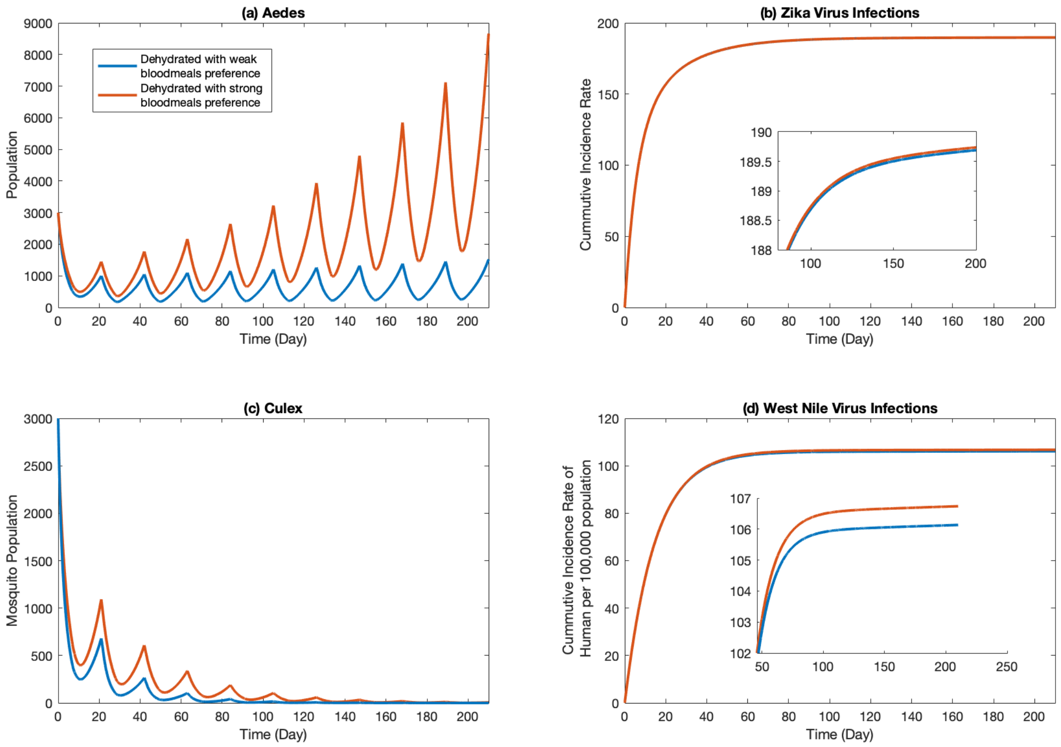


Figure S13: Mosquito population size and related disease scale for two levels of mosquito preference on bloodmeals after repeated cycles alternating between 7 days of dry period followed by various days of wet period (ratio). (a) Population of *Aedes* mosquito; (b) Cumulative number of human infections for Zika virus per 100,000 population; (c) Population of *Culex* mosquito; (d) Cumulative number of human infections for West Nile virus per 100,000 population.

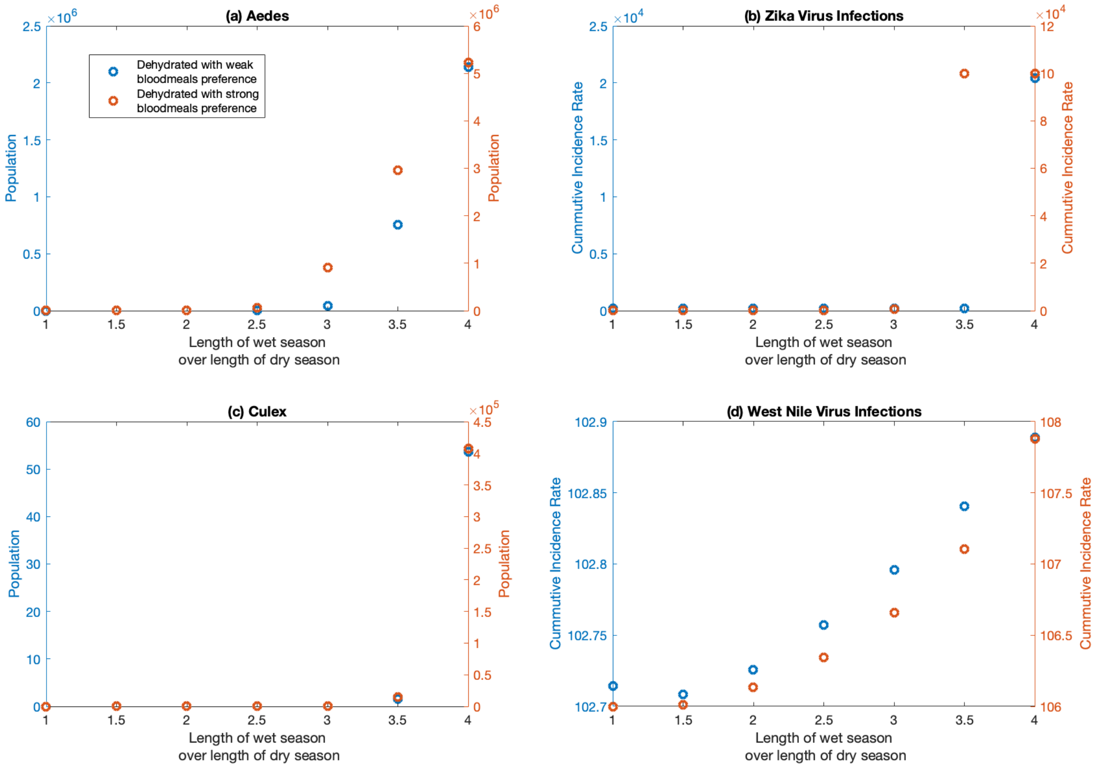


Figure S14: Mosquito population dynamics and related disease dynamics for two levels of mosquito preference on bloodmeals during repeated cycles alternating between 7 days of dry period followed by 14 days of wet period.(a) Population of *Aedes* mosquito; (b) Cumulative number of human infections for Zika virus per 100,000 population; (c) Population of *Culex* mosquito; (d) Cumulative number of human infections for West Nile virus per 100,000 population.
